# Tumor-reactivity of δ2 negative γδT cells and the role of the γδT cell receptor

**DOI:** 10.1101/2021.03.17.435293

**Authors:** Sanne van Dooremalen, Anke Janssen, Lauri Bloemenkamp, Austin Yonika, Dennis X. Beringer, Jürgen Kuball

## Abstract

αβT cells engineered to express a defined γδTCR (TEG) to attack cancer cells have shown great promise when using a γ9δ2TCR to redirect αβT cells (1–9). Reports by us (1–9) and recent reports by others (10, 11) support the key role of the γ9δ2TCR in cancer recognition. We further emphasized the crucial role of the δTCR chain and that differences in CDR3 sequences of the δTCR chain modulates functional avidity of TEGs (2, 8). We and others demonstrated that also δ2 negative γδTCRs are able to redirect αβT cells towards different tumor cell lines (12–15). However, some studies suggest that δ2 negative γδTCRs play a minor role in the tumor recognition by δ2 negative γδT cells (16, 17). In addition for both modes of action for tumor-recognition, δ2 negative γδTCR-dependent and -independent, it has been suggested that CMV infection is not only a major driver of δ2 negative γδT cell expansion (18) but also induces tumor-cross reactive δ2 negative γδT cells (19–21). Therefore, we aimed to systematically explore frequencies of tumor reactive δ2 negative γδT cells in naïve repertoires (cord blood) and patients with or without CMV infection and examined the potential role the parental δ2 negative γδTCR in anti-tumor reactivity of selected clones. We observed that approximately 30% of all tested clones were tumor-reactive, though no differences were observed between different sources. Surprisingly, none of the so far tested γδTCR did mediate strong anti-tumor reactivity of the parental clones. Though numbers of tested TCR sequences are still low, our data imply that tumor-reactivity of δ2 negative γδT cells is frequently not mediated by the δ2 negative γδTCR alone.

## Material and Methods

### Isolation and sorting of CBMC/PBMCs

CBMC/PBMCs from donor blood were isolated with a Ficoll-Paque density gradient and stained with a Vδ2-FACS panel, consisting of the following fluorochome labeled antibodies; CD3-PB (1:20; clone UCHT1; Beckman Coulter), CD4-PE-Cy7 (1:100; RPA-T4; eBiosciences), CD8-PE-Cy5.5 (1:1000; RPA-T8; biolegend), pan-γδTCR-APC (1:5; B1; BD Biosciences), Vδ1-FITC (1:10; TS8.2; Thermo scientific), Vδ2-PE (1:25; B6; BD Biosciences), in RPMI-1640 supplemented with 10% human serum, 1% Penicillin/Streptomycin and 25 nM β-mercaptoehtanol (=10% huRPMI). Using a BD FACSAria II, CD3+γδTCR+ Vδ2-cells were sorted at 100 cells per well into 96 wells U-bottom plates, containing 200 μl 10% huRPMI and “γδ Rapid Expansion Protocol mix”, consisting of irradiated feeder cells, IL-2, IL-15 and PHA-L. The sorted oligoclones were maintained on several REP cycles to obtain sufficient T cells for functional analysis, TCR sequencing, cryopreservation in 90% human serum + 10% DMSO, and/or isolation of single cell clones.

### Tumor panel cell lines

Daudi, ML-1, K562, and Jurkat-76 were cultured in RPMI1640 supplemented with 10% FCS and 1% Penicillin/Streptomycin and passaged twice a week. MDA-MB-157, MDA-MB-231, MZ1851RC, OVCAR-3, SCC-9, and HT-29 cells were cultured in DMEM supplemented with 10% FCS and 1% Penicillin/Streptomycin. These adherent cells were passaged twice a week, using trypsin-EDTA to detach the cells.

### HLA-typing

Cell linesMDA-MB-157, MDA-MB-231, MZ1851RC, OVCAR-3, SCC-9, and HT-29 and T cells of PBMC donors DS81, DS82, and DS83 were send for HLA phenotype analysis at the HLA-laboratory of the UMCU.

### INFγ ELISA

The IFNγ production of Vδ2-oligoclones against the panel of tumor cells was tested with the enzyme-linked immunosorbent assay (ELISA). The oligoclones and tumor cells were counted and 100k effectors and 100k target cells were added together in a 96-wells plate in a final volume of 200 μL. T cells transduced with LM1, and Clone 5 were taken along as controls. The cells were incubated overnight and IFNγ levels in the supernatant were detected using the human IFN-γ ELISA Ready-SET-Go! kit (Invitrogen) accorded to manufacturer’s instruction.

### Isolation of single cell clones

Reactive oligoclones derived from cord blood were isolated to obtain single cell clones. Using the same FACS panel as stated above several oligoclones were single cell sorted using FACS. Again the single cells were expanded using the γδT cells REP protocol.

For cord blood donor 4 (CB-4), the most reactive oligoclones were selected for limiting dilution. Cells were diluted to 3.3 cells/ml and plated in a 96 wells plate at 100 μL/well, 100 μL of γδREP mix were added to the wells and expanded using the γδT cells REP protocol.

### RNA isolation and synthesizing, purification and amplification of cDNA and HTS sequencing

RNA of reactive oligoclones (300.000 cells) was isolated using the RNease minikit (Quiagen) according to manufacturer’s instruction. cDNA was synthesized from the isolated RNA by using the superscript II reverse transcriptase kit (Invitrogen) according to manufacturer’s instructions in combination with gamma and delta specific primers, followed by a purification step using the PCR clean-up Kit (Machery-Nagel) accorded to manufacturer’s instruction.

The cDNA was amplified with PCR using Phusion polymerase. Thereafter, the samples were ran on a 1% agarose gel And the DNA was purified with the gel clean-up Kit (Machery-Nagel) accorded to manufacturer’s instruction. The concentration of cDNA was measured with a nanodrop and the samples were sent for high throughput sequencing as described previously (2, 14). HTS data in the standard FASTQ format was processed using MiXCR (version-v2.1.1) (22)for TCR sequence alignment, assembly and clonotype extraction.

### Cloning matched TCR sequences in pBullet

The TCR sequences of selected oligoclones were generated using overlap extension PCR where the CDR3 sequences of existing TCR clones were substituted with the frequency matched oligoclone sequences as previously described for γ9δ2 TCRs (2). The new sequences were subcloned in pBullet using NcoI and BamHI restriction sites.

### Transduction of OC-TCRs in Jurkat-76

Retroviral vector pairs of oligoclones were used to transduce Jurkat-76 cells as decribed previously. In short, pBullet-OC-TCRγ and pBullet-OC-TCRδ together with pHit60 and pColtGALV were incubated with Fugene and used to transfect PhoenixAmpho cells. Virus containing supernatants of these PhoenixAmpho cultures, supplemented with polybrene, were used to transduce Jurkat-76 cells. Antibiotics geneticin and puromycin were added to the culture to select for successfully transduced Jurkat-76.

### Functional Testing of OC-TCRs in transduced Jurkat-76 cells

Functional testing of OC TCRs expressed in Jurkat-76 was done by measuring CD69 upregulation after an overnight co-culture of 10^5^ effector cells (EC) with indicated target cell lines (TC) at 1:1 ratio. Prior to co-culturing the target cells were labeled using Cell Trace Violet (1:5000; Thermofisher). Cells were stained with following antibody mix for 30 minutes on ice: pan-γδTCR-PE (1:10; IMMU510; Beckman Coultier) and CD69-APC (1:25; FN50; Sony Biotechnology). The samples were analyzed using a BD FACSCanto II (BD Bioscience) and the readouts were analyzed using BD FACS Diva software.

## Results

### Characterization of tumor reactive δ2^neg^ T cell repertoires

The potential variety of the γδTCR repertoire has been suggested to exceed by 2–3 log the theoretical diversities of antibodies and αβTCR (23). Therefore, our screening strategy to assess frequency of tumor reactive γδT cell repertoire and the subsequent isolation of potentially tumor-reactive γδTCR was based on creating first oligoclonal cultures through sorting and expanding 100 δ2 negative γδT cells per well followed by functional testing of these oligoclones. Three different sources of donors were used. Cord blood from 4 donors was used as source for naïve immune repertoires with many invariant γδTCR chains (24). PBMCs from 3 CMV-negative and 3 CMV-positive healthy donors were used in order to assess the impact of CMV infection on the frequency of tumor reactive γδT cells during adolescence. We hypothesized in line with previous observations (20, 25) that higher numbers of δ2 negative γδT cells can preferentially be found in donors which had a CMV infection. FACS-based sorting included antibody panel directed against CD3, CD4, CD8, panγδTCR, δ1TCR, and δ2TCR. The percentage of γδTCR positive cells varied between 0.6-7.0% of the CD3^+^ T cells. There was a large variation in the percentage of δ2 negative γδT cells among total number of γδT cells. As expected, the majority of γδT cells derived from cord blood donors were δ2 negative γδT cells (73.3 to 91.7%) (26, 27), while CMV negative adult donors had the lowest numbers (34.5 to 45%) and CMV positive donors had the widest range (43.4 to 100%) (**Supplementary Figure1**).

In order to assess potential tumor reactivity of different immune repertoires against tumors, up to 96 oligoclones were sorted per donor (Table 1) and expanded using several rounds of the rapid expansion protocol, as described in materials and methods, to have sufficient cells for assessing their anti-tumor reactivity. In order to test anti-tumor reactivity of the oligoclones, we selected a panel of tumor cell lines that have been used partially also for screening of tumor reactivity of TEG001 (2,3,5,6,8,9,14). Cell lines included hematological tumor cell lines, Daudi, ML-1, and K562, and the solid tumor cell lines, MDA-MB-157, MDA-MB-231, MZ1851RC, OVCAR-3, SCC-9, and HT-29. An ELISA was used to determine the IFNγ concentration in the media after co-culture of T cells with tumor cells. A randomly set threshold of IFNγ concentration of at least 15 pg/ml and at least 1.5 times higher IFNγ concentration compared to the oligoclone alone was used to define reactivity within the oligoclonal pool. 215 of the 768 tested oligoclones were reactive to one or more cell lines (**Table 1** and **Supplementary Figure 1**), while 39 clones were hyperactive already in the absence of a target. Most tumor reactive oligoclones recognized only one (n=106) or two cell lines (n=58), while 50 oligoclones recognized more than two cell lines (**Table 2**). There were no major differences between the cord blood and peripheral blood derived oligoclones in their ability to recognize multiple cell lines. INFγ secretion was for the majority of the reactive oligoclones below 100 pg/ml. Only a few produced high levels of IFNγ (>500 pg/ml) and in most cases these cell lines were only reactive towards one tumor cell line. The more broadly reactive oligoclones secreted lower levels of IFNγ (**Supplementary Figure 2**). We furthermore typed all tumor cells if applicable for HLA class I in order to get first hints for potential alloreactive HLA-restricted γδT TCR based on recognition patterns of clones, as reported (15) (Supplementary Table 2). However, none of the multiple tumor-reactive clones followed a defined HLA pattern.

**Table 1.** Summary of reactivity against the tumor panel cell lines of peripheral blood derived oligoclones.

| Donor | CMV status | Tested<br>(n) | Tumor-reactive<br>(n) | Hyperactive<br>(n) | Tumor-reactive<br>(%) | Hyperactive<br>(%) |
| --- | --- | --- | --- | --- | --- | --- |
| CB-1 | n.t. | 27 | 7 | 2 | 25.9 | 7.4 |
| CB-2 | n.t. | 30 | 5 | 4 | 16.7 | 13.3 |
| CB-3 | n.t. | 96 | 35 | 0 | 36.5 | 0 |
| CB-4 | n.t. | 53 | 15 | 6 | 28.3 | 11.3 |
| **Total** |  | **206** | **62** | **12** | **30.1** | **5.8** |
| PB-DS67 | Neg | 82 | 1 | 2 | 1.2 | 2.4 |
| PB-DS81 | Neg | 96 | 32 | 7 | 33.3 | 7.3 |
| PB-DS82 | Neg | 96 | 36 | 6 | 37.5 | 6.3 |
| **Total** | **Neg** | **274** | **69** | **15** | **25.2** | **5.5** |
| PB-DS71 | Pos | 96 | 6 | 1 | 6.3 | 1.0 |
| PB-DS80 | Pos | 96 | 36 | 8 | 37.5 | 8.3 |
| PB-DS83 | Pos | 96 | 42 | 3 | 43.8 | 3.1 |
| **Total** | **Pos** | **288** | **84** | **12** | **29.2** | **4.2** |
| **Total** |  | **768** | **215** | **39** | **28.1** | **5.2** |

**Table 2.** overview of the number of cell lines recognized by the oligoclones.

| No. Targets recognized | 1/8 | 2/8 | 3/8 | 4/8 | 5/8 | 6/8 | 7/8 | 8/8 |
| --- | --- | --- | --- | --- | --- | --- | --- | --- |
| CB-Oligoclones (No.) | 30 | 19 | 8 | 2 | 2 | 0 | 1 | 0 |
| CB-Oligoclones (% of all) | 14.6 | 9.2 | 3.9 | 1.0 | 1.0 | 0.0 | 0.5 | 0.0 |
| PB-Oligoclones (No.) | 76 | 39 | 20 | 3 | 9 | 6 | 0 | 0 |
| PB-Oligoclones (% of all) | 13.5 | 6.9 | 3.6 | 0.5 | 1.6 | 1.1 | 0.0 | 0.0 |
| **Total (No.)** | **106** | **58** | **28** | **5** | **11** | **6** | **0** | **0** |
| **Total ((% of all)** | **13.8** | **7.6** | **3.6** | **0.7** | **1.4** | **0.8** | **0** | **0** |

### High throughput sequencing of reactive oligoclones to determine TCR sequences by frequency matching

Initially, single cell cloning of oligoclones was the preferred strategy to obtain matched TCRγ and TCRδ sequences. However, the expansion and survival rate of single cell sorted oligoclones was very low. After several fruitless attempts on the cord blood samples, resulting in only 2 TCR sequences (Supplementary Table 2), an alternative strategy was used to obtain the TCR sequences. We used our established pipeline for high throughput sequencing (HTS) of TCR repertoires to determine the relative frequencies of TCRγ and TCRδ sequences. Depending on the relative abundance of the TCRγ and TCRδ chains an educated guess can be made which sequences to pair. Selection of oligoclones for further sequencing was based either on broader tumor-reactivity, defined as recognizing 5 or 6 cell lines (n= 8), or high potency, defined as >500 pg/ml IFNγ (n= 5). In total 20 oligoclones were selected for repertoire sequencing and of 16 oligoclones both the TCRγ and TCRδ chains were successfully sequenced. For the other 4 oligoclones insufficient cDNA obtained for sequencing. For the majority of oligoclones one TCRδ sequence and one TCRγ sequence were most prevalent and were selected for further cloning in the pBullet retroviral expression vectors. While for other oligoclones, the combination was less evident (**Figure 1**). For three of the oligoclones, OC6, OC100, and OC180, we selected one TCRδ sequence and two TCRγ sequences, for OC151 we selected two TCRδ sequences and one TCRγ sequence, and for OC47 we selected two TCRδ sequences and two TCRγ sequences for further cloning (Supplementary Table 2). To identify the correct TCRδ and TCRγ sequences, combinations of these TCR chains need to be tested for functionality in separate transductions of T cells.

**Figure 1.**
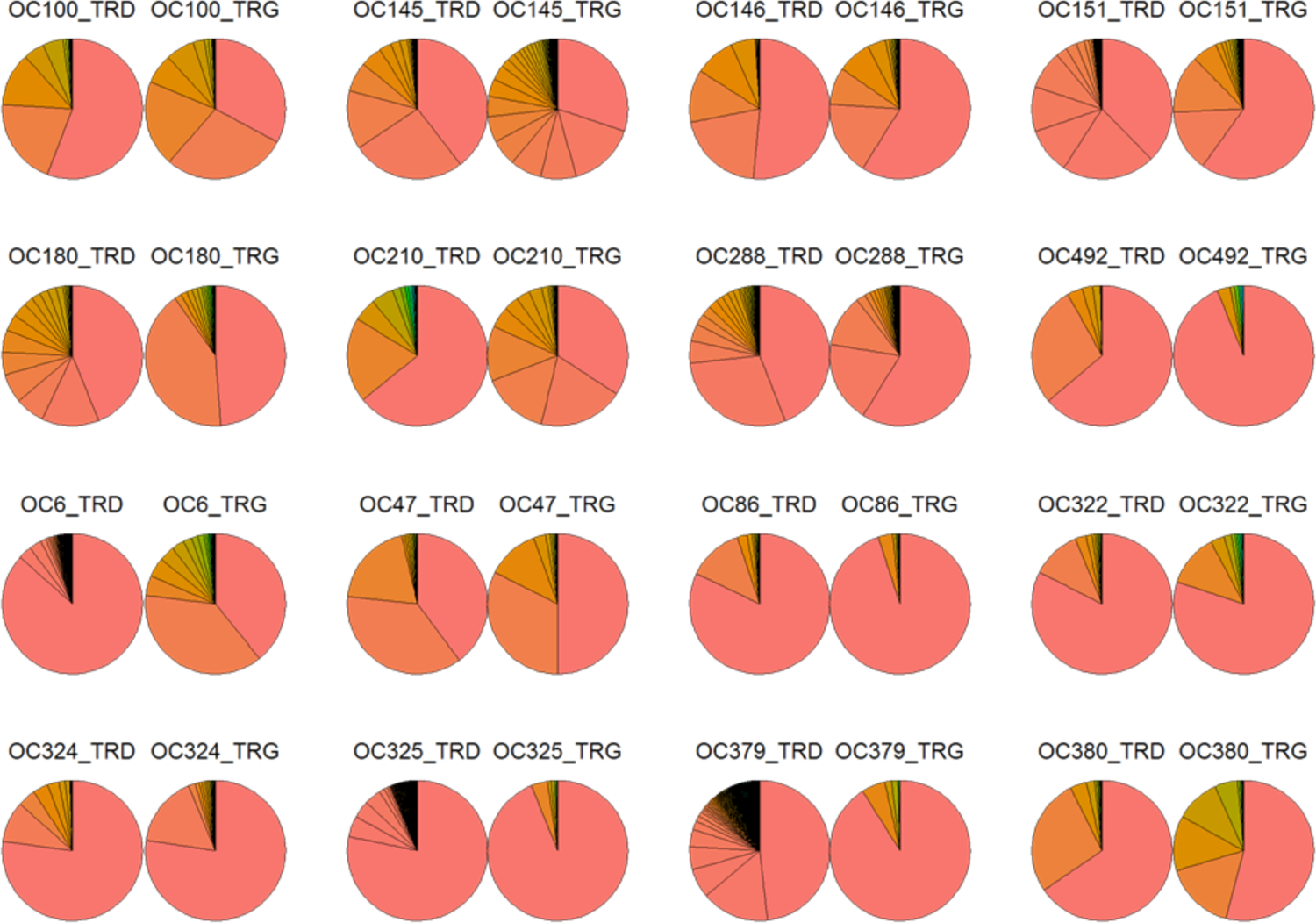
clonal frequencies of TCRδ and TCRγ sequences within the oligoclones. The pie plots represent the frequency of a TCRδ (_TRD) sequence or TCRγ (_TRG) sequence within an oligoclone based on the number of sequence reads in the HTS results. Based on frequency matching the one or two sequences were used for further cloning.

### Tested combinations did not show reactivity towards cell lines as their parental clones

For further analyses oligoclones OC47 and OC288 were selected for testing as these two oligoclones showed high reactivity (INFγ >500 pg/ml) to one of the tumor cell lines in the panel. Moreover, OC288 had one prevalent TCRδ and one prevalent TCRγ sequence in the HTS results, while OC47 had two TCRδ and two TCRγ sequences with comparable prevalence in the repertoire. We used Jurkat-76 (J76) cells transduced with the selected oligoclone derived TCRs, OC288 and four combinations of OC47. In addition, J76 were transduced with controls, γ9δ2TCRs LM1 as negative control and Cl5 as positive control. After antibiotic selection the transduced J76 cells were analyzed by FACS for TCR expression. The two γ9δ2TCRs LM1 and Cl5 and the δ2neg TCRs clone OC47ba had comparable expression. The other oligoclone derived TCRs showed a lower surface expression (**Figure 2A**). We used the same tumor cell panel as was used for testing the oligoclones to assess the specificity of the transduced J76 cells. Tumor cells were incubated with the transduced J76 cells overnight in a 1:1 effector-to-target ratio, activation of the T cells was assessed by measuring CD69 surface expression using FACS. The two reference TCRs showed the expected results, no reactivity for J76-LM1 towards any of the cell lines, while J76-Cl5 were activated by Daudi and MZ1851RC, the extent of activation of the J76-Cl5 was enhanced in presence of 100 µM pamidronate (**Figure 2B**). J76 cells transduced with one of the oligoclones derived TCRs showed hardly any activation. OC288 produced high levels of IFNγ when it was incubated with Daudi, but the reconstructed TCR transduced in J76 did not show any activation in the presence of Daudi. Likewise, OC47 was strongly activated by OVCAR-3 cells, however none of the four combinations showed reactivity towards this tumor cell line (**Figure 2B**). When we normalized the data, to be able compare the different J76-transductants, we saw that some of the δ2 negative TCR showed a slight increase in CD69 expression, e.g. J76-OC47aa had increased surface expression when incubated with MDA-MB-231 and HT-29, but none of them exceeded the 1.5 fold change line (**Supplementary Figure 3**).

**Figure 2.**
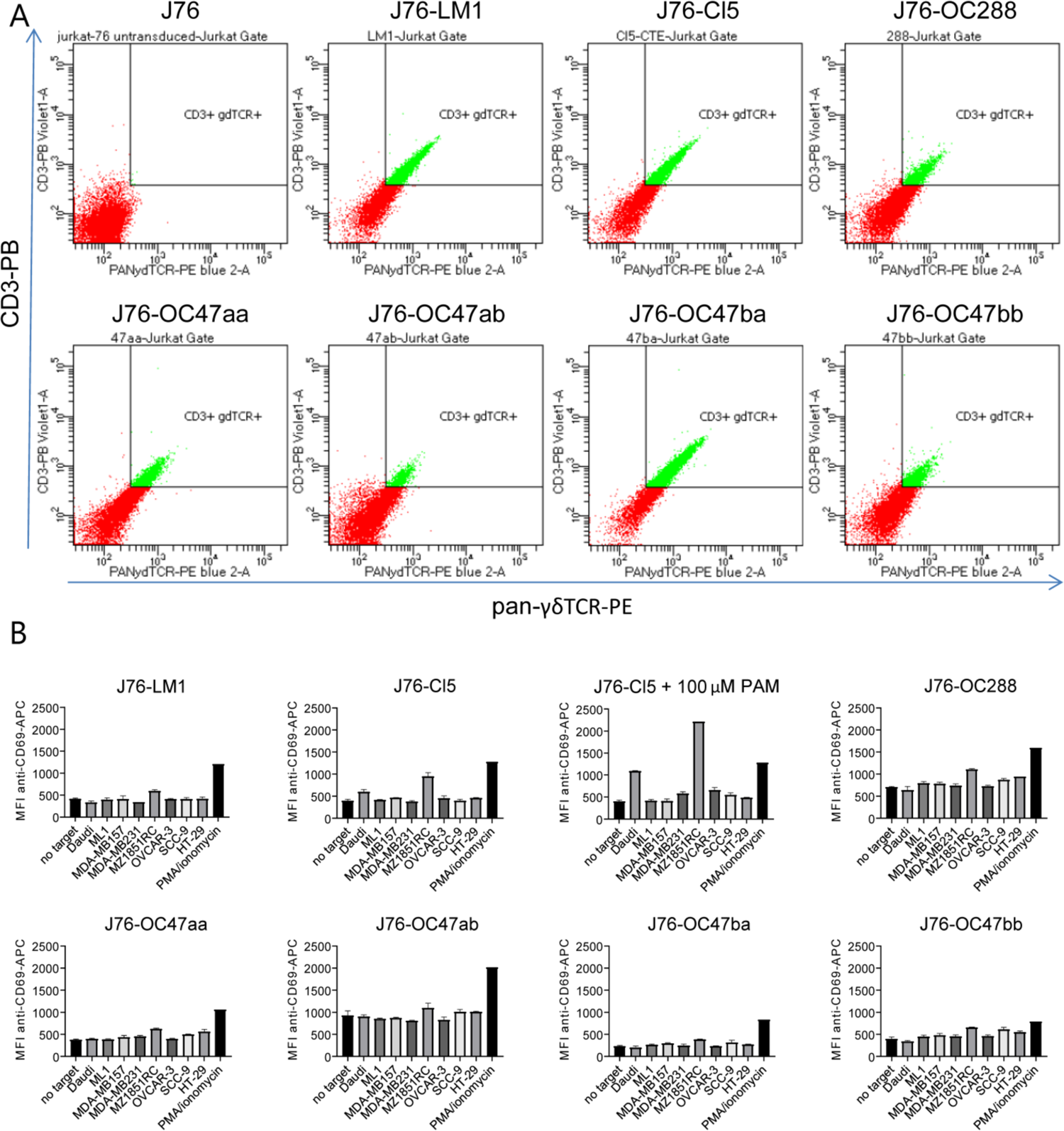
TCRs derived from tumor reactive oligoclones do not show reactivity when transduced in Jurkat-76 cells. (A) Surface expression of the transduced γδTCR was analyzed by FACS. Cells were stained with fluorescently labeled antibodies against CD3 (pacific blue) and pan γδTCR (PE). (B) transduced Jurkat-76 cells were incubated with target cells labeled with cell trace violet for 16 h. The cells were stained for surface expression of CD69, CD3, and γδTCR, and analyzed by FACS. Plotted are the mean fluorescent intensity (MFI) values for CD69-APC on the γδ+ Jurkat-76 cells.

## Discussion

Main finding of our study is that regardless of the used immune repertoire we found that approximately one third of generated δ2 negative γδT cells are reactive against different tumor cell lines. However, reactivity was rather scattered in terms of number and types of tumors being recognized. We found “super spreaders” which have been able to recognize many different tumor types while other clones had a very focused and strong reactivity against a single tumor when assessed by IFNγ secretion. A major technical challenge for the analyses of cord blood derived oligoclones was the limited ability for expansion. Single cell sorting of reactive oligoclones from cord blood did not result in sufficient numbers of cells to assess their reactivity towards extended tumor cell panels. Many of the single cell clones did not grow out at all, others could not be expanded beyond two or three cycles, making later analyses impossible. This observation was made for four different donors suggesting that the nature of cord blood derived clones is substantially different from clones generated from the adult immune repertoire (16,24,28,29) which can be expanded over many cycles (14). In contrast, larger number of sequences were obtained using high throughput sequencing of 16 analyzed reactive oligoclones from adults. Though many of the oligoclones had one prevalent TCR sequence, some had more. Given these major technical hurdles, to date only limited numbers of γδTCRs have been cloned and functionally tested. Selected TCR pairs which were tested in Jurkat-76 cells for reactivity did not show any reactivity towards any of the cell lines in the panel, implying that these TCRs are at least alone not tumor reactive. Whether the other identified TCRs are tumor reactive remains to be determined.

---

**Supplementary Figure 1.**
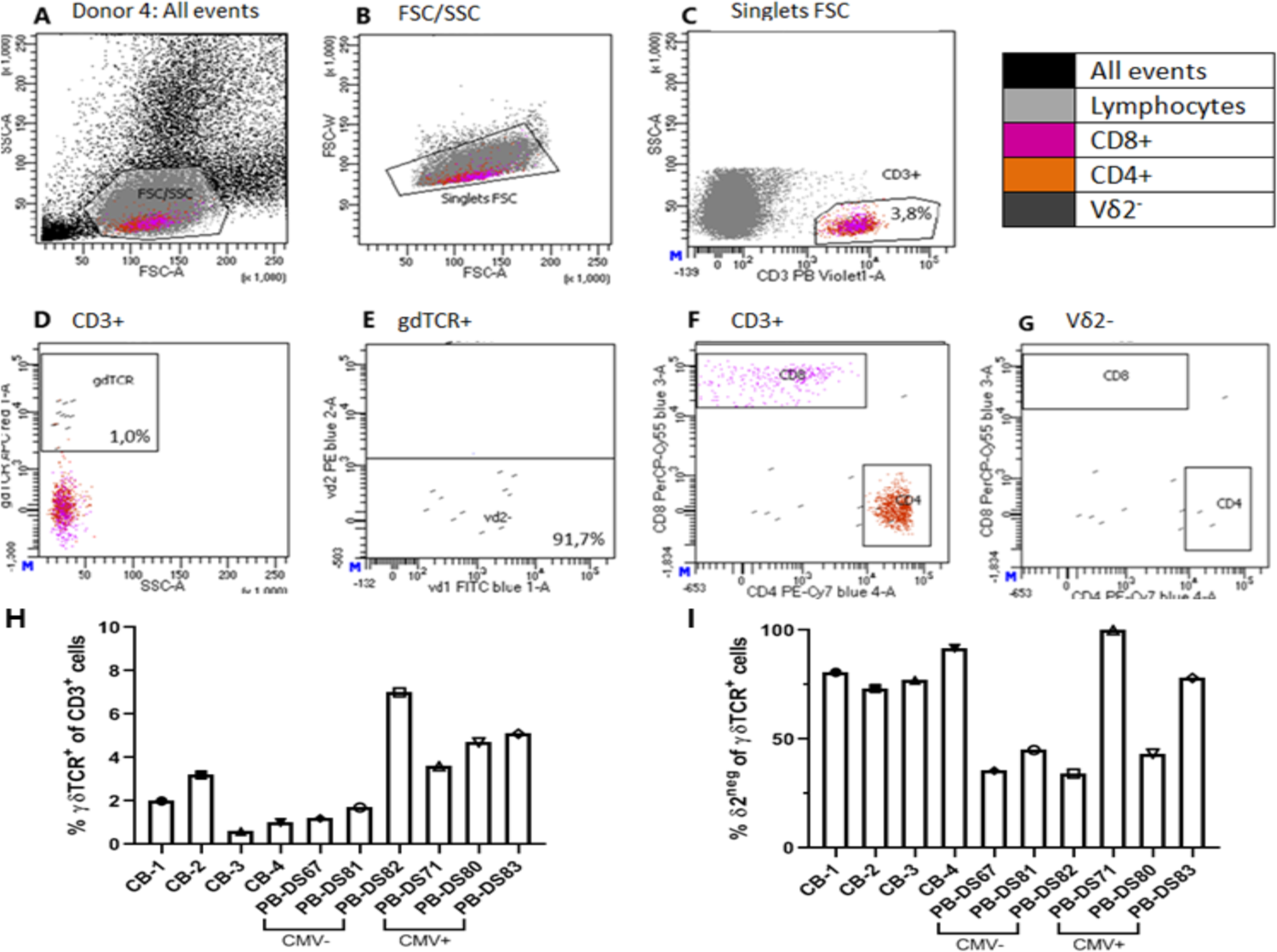
(A-G) Representative gating strategy for sort of δ2 TCR negative cells from donor CB-4. (H) percentage of γδTCRpos cells in sorted samples. (I) percentage of δ2-TCR negative cells in sorted samples

**Supplementary Figure 2.**
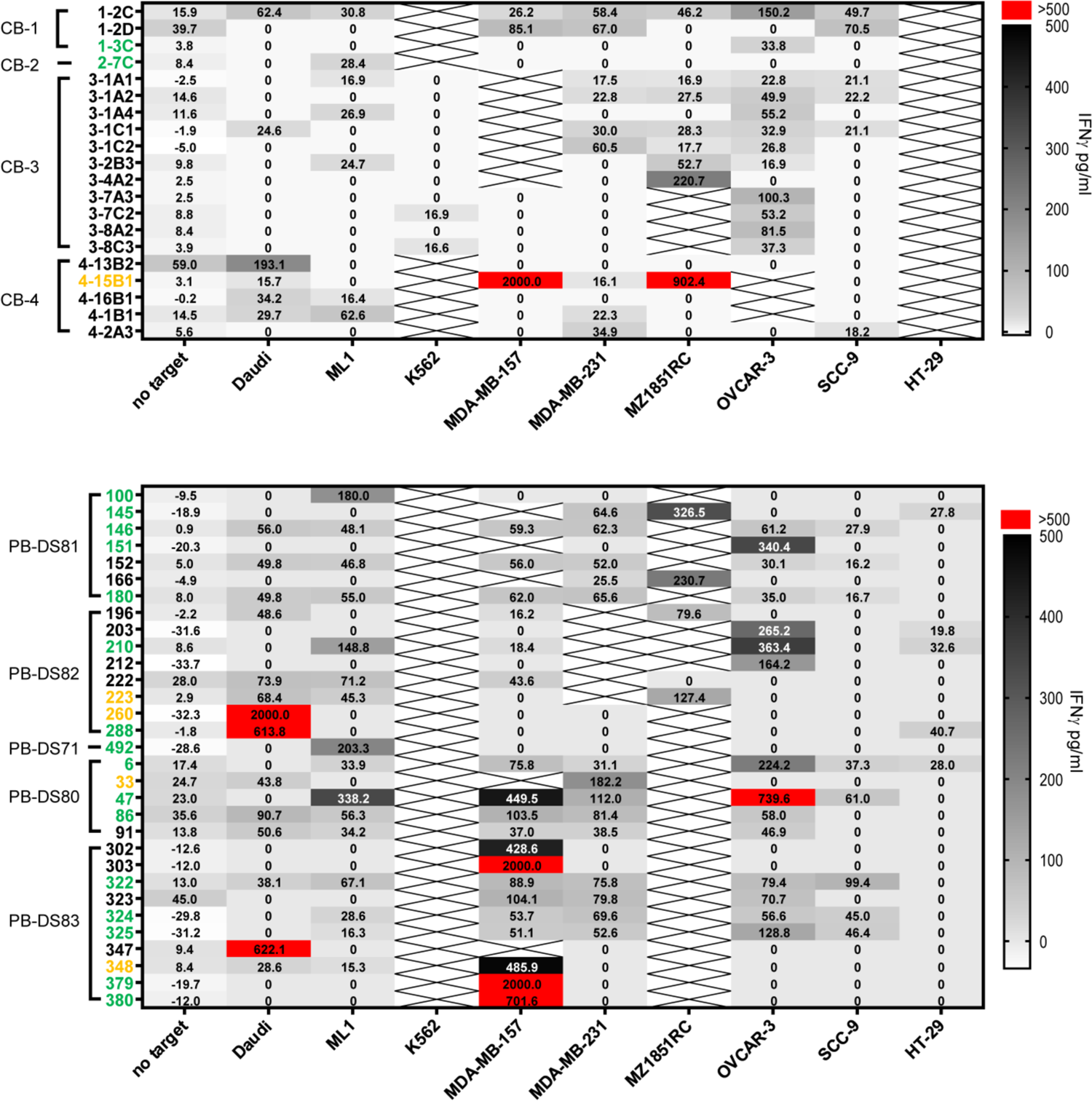
Oligoclone reactivity towards tumor cell line panel of the most reactive clones. 10^5 δ2^neg^ T cells were incubated in a 1:1 ratio with the indicated tumor cell lines for 16-20 hours. INFγ production was detected using an ELISA and used a marker for T cell activation. Oligoclones numbers indicated in green have been successfully sequenced, no sequences were obtained for those colored orange. The data shown is modified to accentuate the reactivity, non-reactive condition have the arbitrary value of 0 pg/ml. Appendix 1 contains non-modified data for each donor.

**Supplementary Figure 3.**
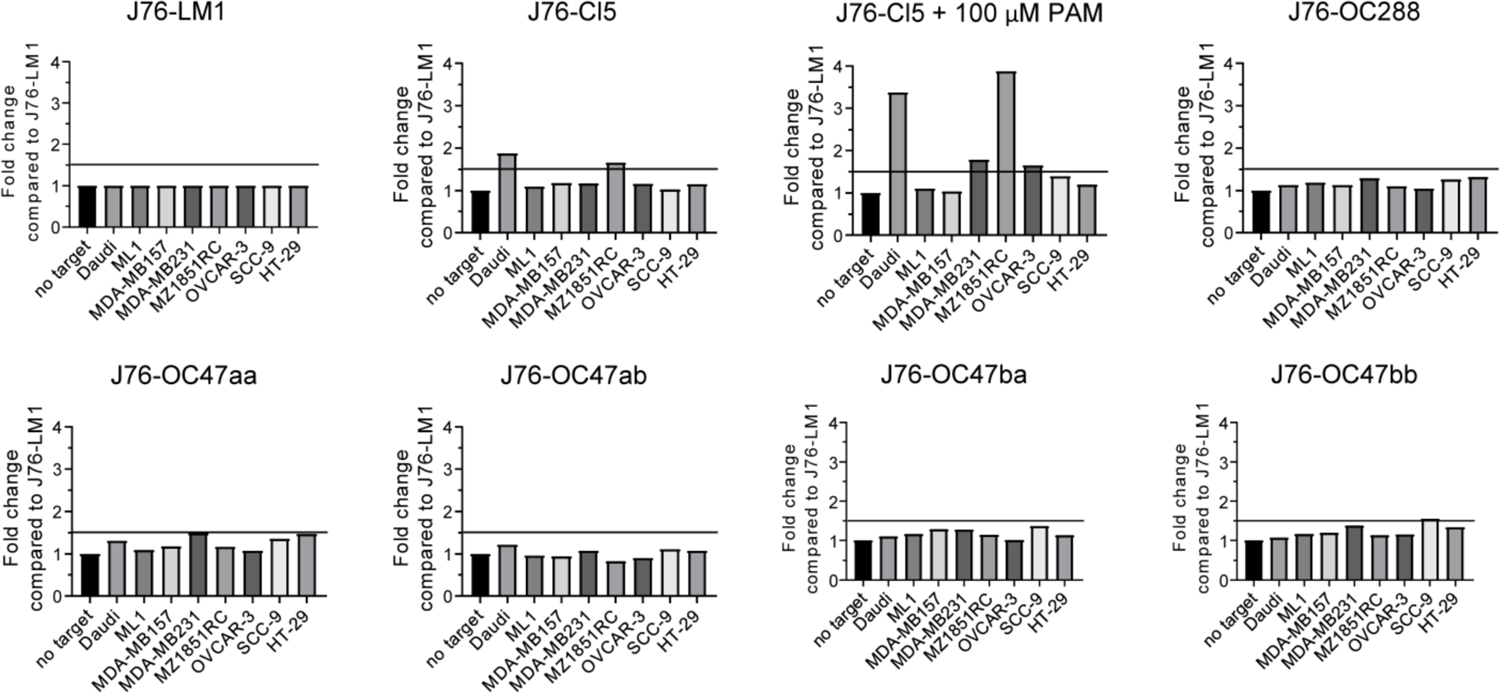
TCRs derived from tumor reactive oligoclones do not show reactivity when transduced in Jurkat-76 cells. Normalized CD69 surface expression data from co-incubation assay. First, all conditions of a individual TCR were normalized to the CD69 expression of J76-TCR without target. Next, within the stimulation conditions all different TCRs were normalized to the already normalized J76-LM1 condition. Related to figure 2.

**Supplementary Table 1.**
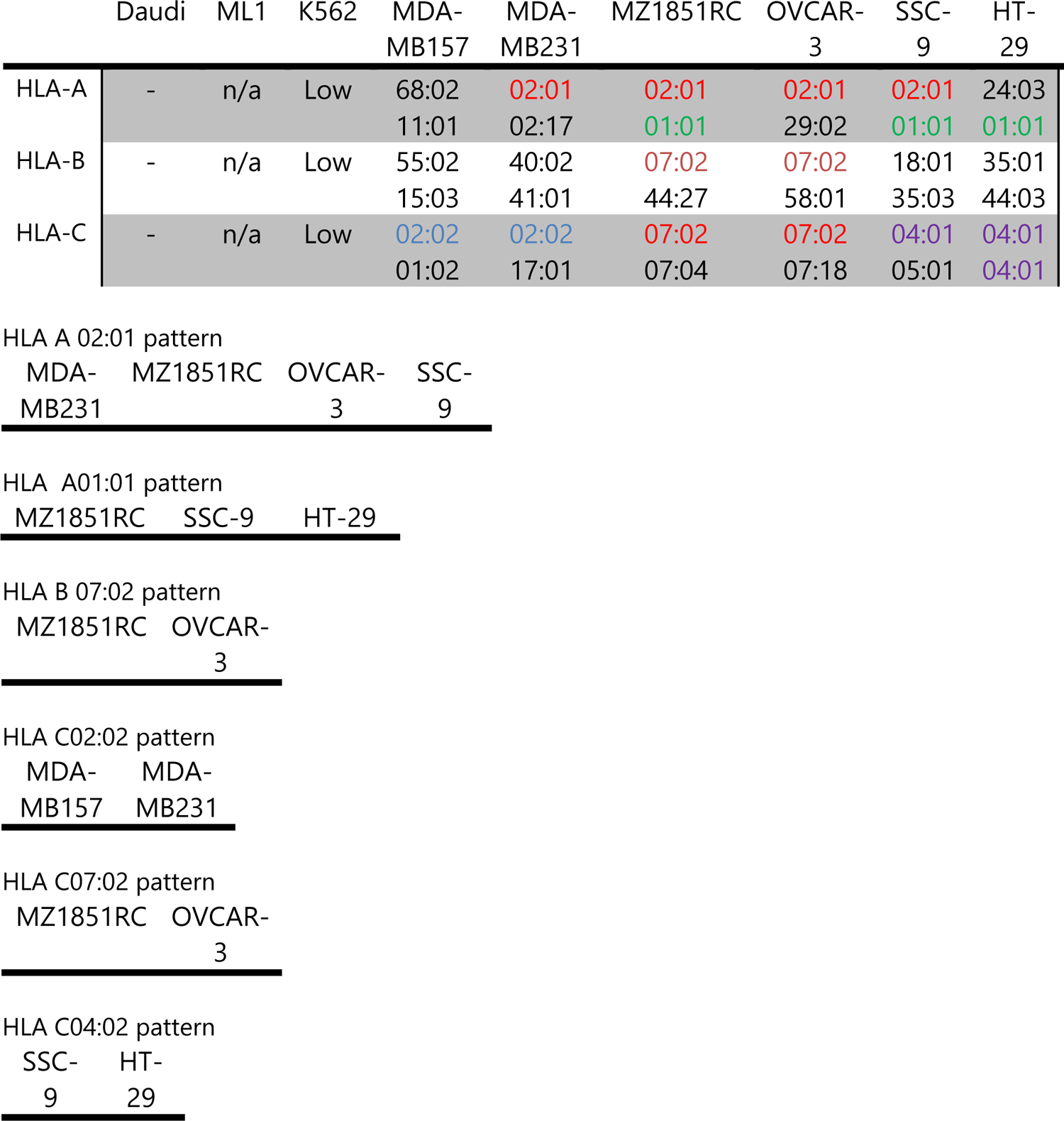
HLA-class I phenotypes of cell lines

|  | Daudi | ML1 | K562 | MDA-MB157 | MDA-MB231 | MZ1851RC | OVCAR-3 | SSC-9 | HT-29 |
| --- | --- | --- | --- | --- | --- | --- | --- | --- | --- |
| HLA-A | - | n/a | Low | 68:02<br>11:01 | 02:01<br>02:17 | 02:01<br>01:01 | 02:01<br>29:02 | 02:01<br>01:01 | 24:03<br>01:01 |
| HLA-B | - | n/a | Low | 55:02<br>15:03 | 40:02<br>41:01 | 07:02<br>44:27 | 07:02<br>58:01 | 18:01<br>35:03 | 35:01<br>44:03 |
| HLA-C | - | n/a | Low | 02:02<br>01:02 | 02:02<br>17:01 | 07:02<br>07:04 | 07:02<br>07:18 | 04:01<br>05:01 | 04:01<br>04:01 |

HLA A 02:01 pattern
|  |  |  |  |
| --- | --- | --- | --- |
| MDA-MB231 | MZ1851RC | OVCAR-3 | SSC-9 |

HLA A01:01 pattern
|  |  |  |
| --- | --- | --- |
| MZ1851RC | SSC-9 | HT-29 |

HLA B 07:02 pattern
|  |  |
| --- | --- |
| MZ1851RC | OVCAR-3 |

HLA C02:02 pattern
|  |  |
| --- | --- |
| MDA-MB157 | MDA-MB231 |

HLA C07:02 pattern
|  |  |
| --- | --- |
| MZ1851RC | OVCAR-3 |

HLA C04:02 pattern
|  |  |
| --- | --- |
| SSC-9 | HT-29 |

**Supplementary Table 2.**
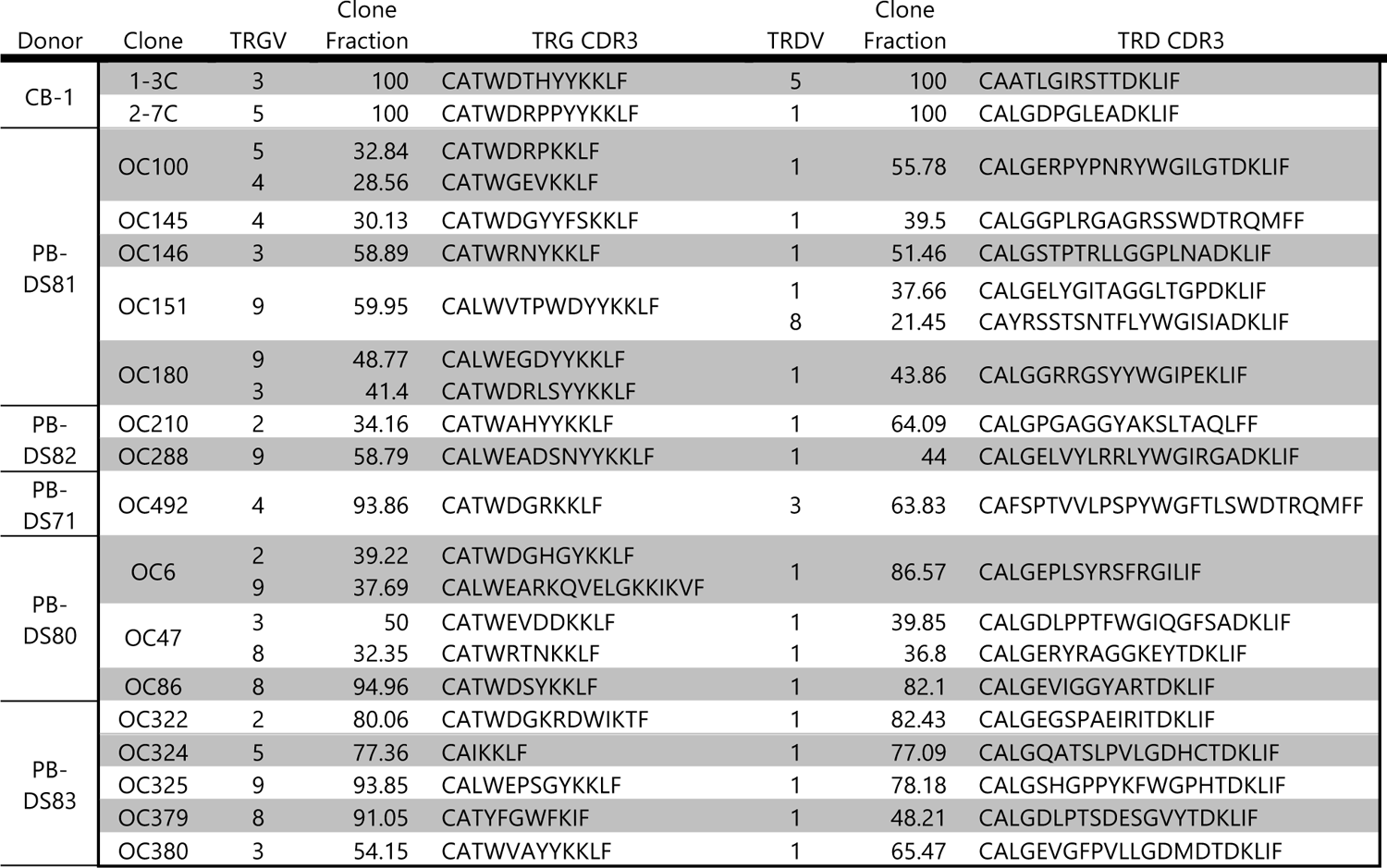
Overview TCR gene usage and CDR3 sequences of sequenced δ2 negative TCRs

## Appendix

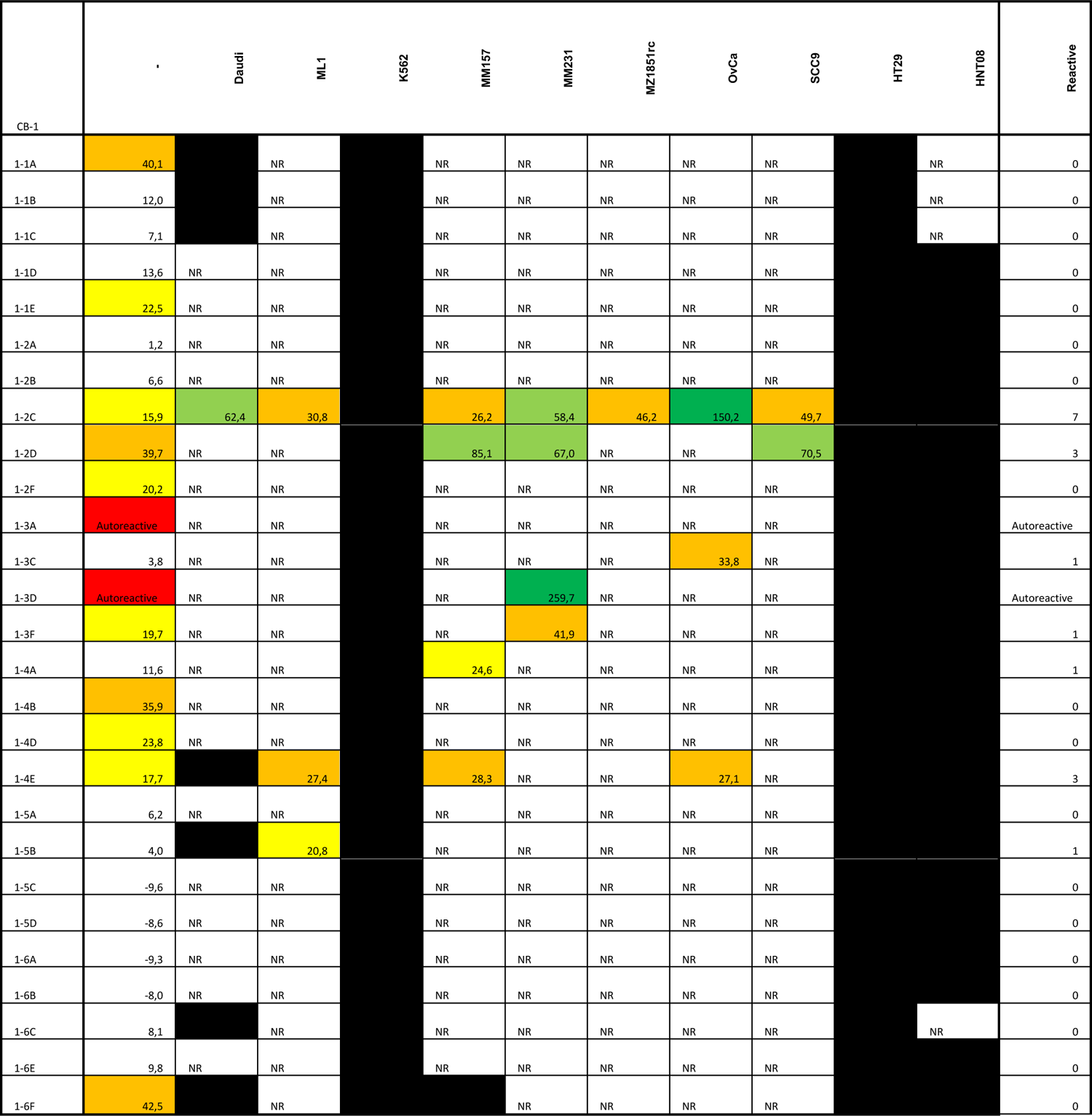
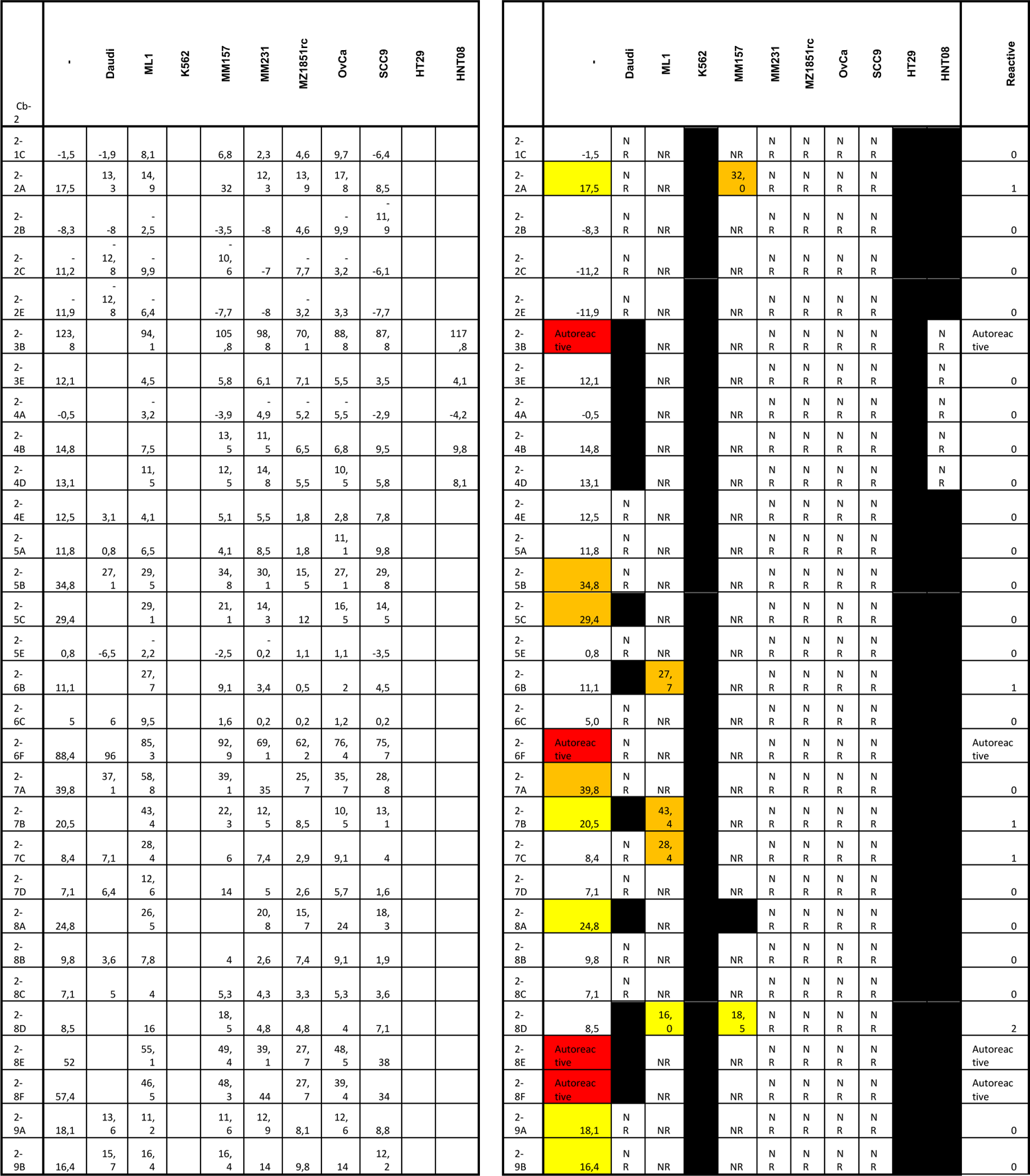
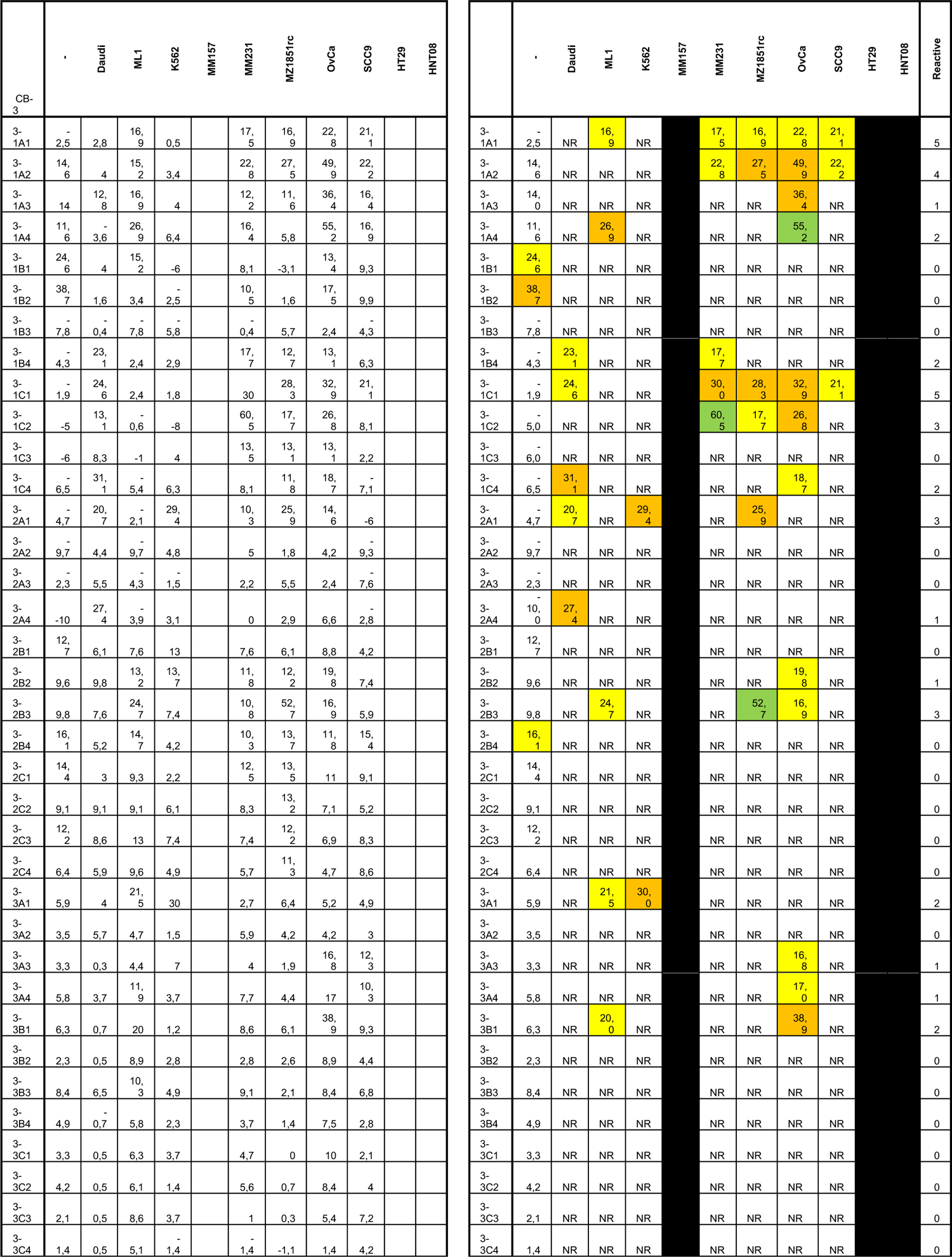
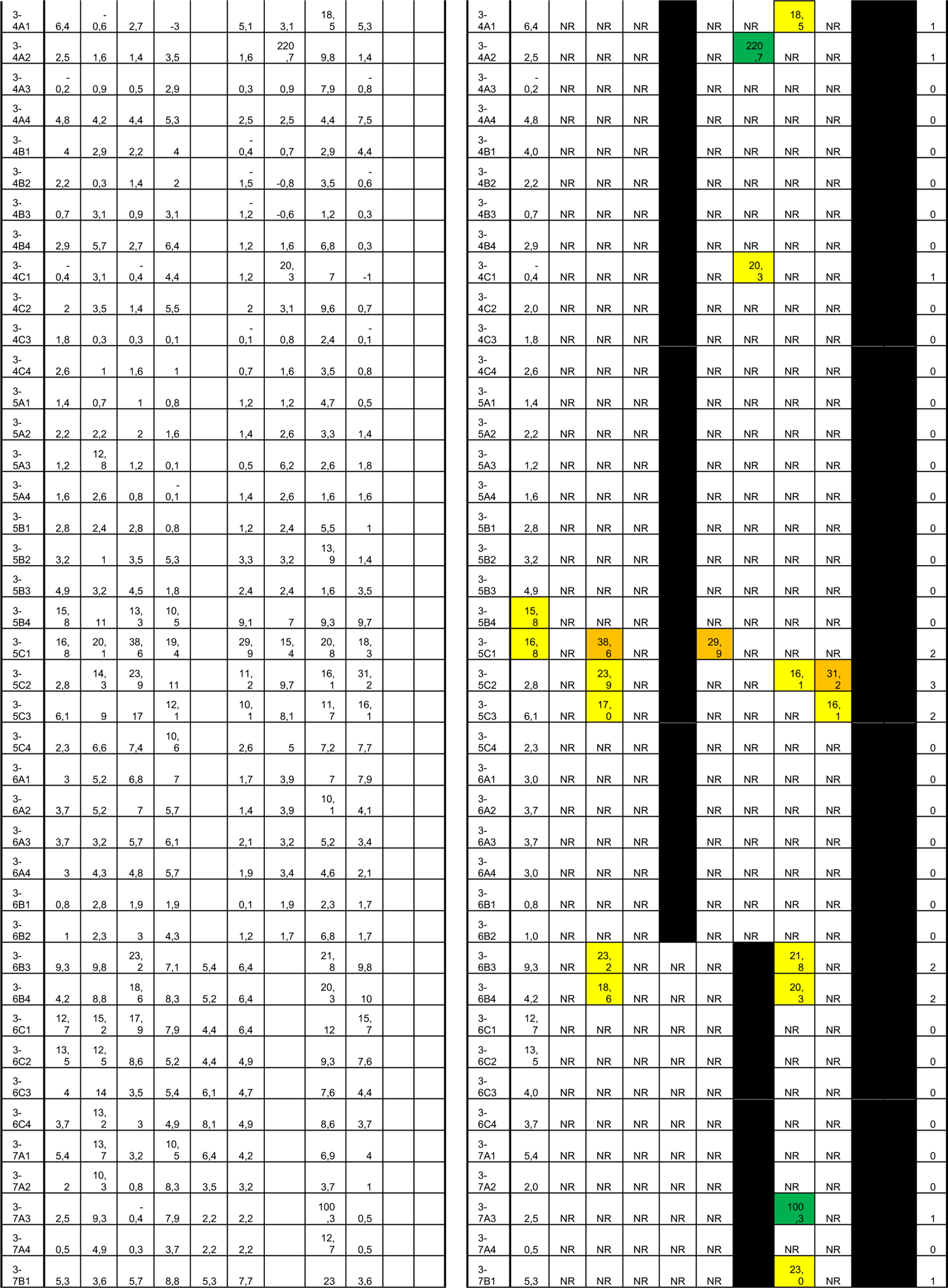
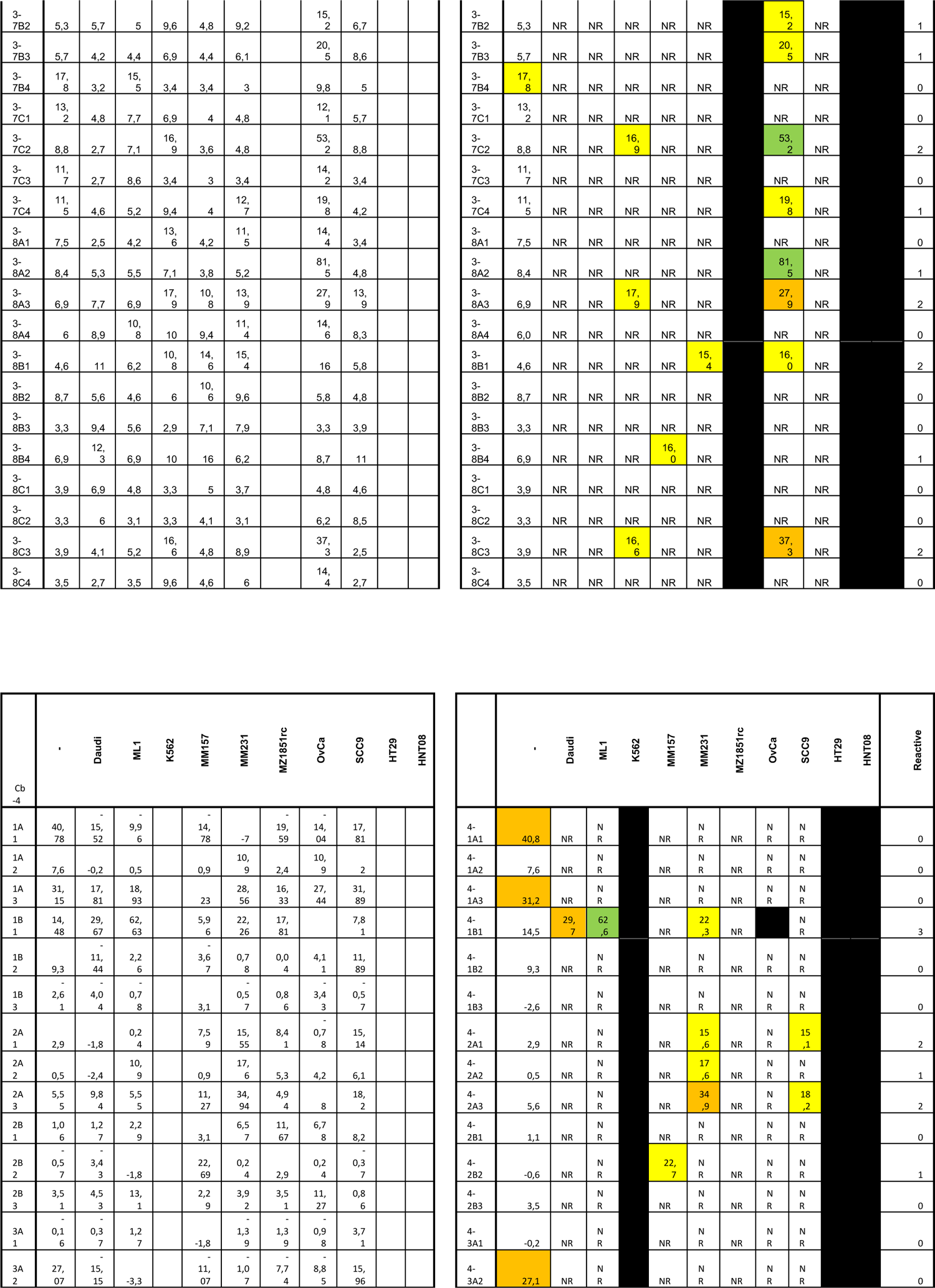
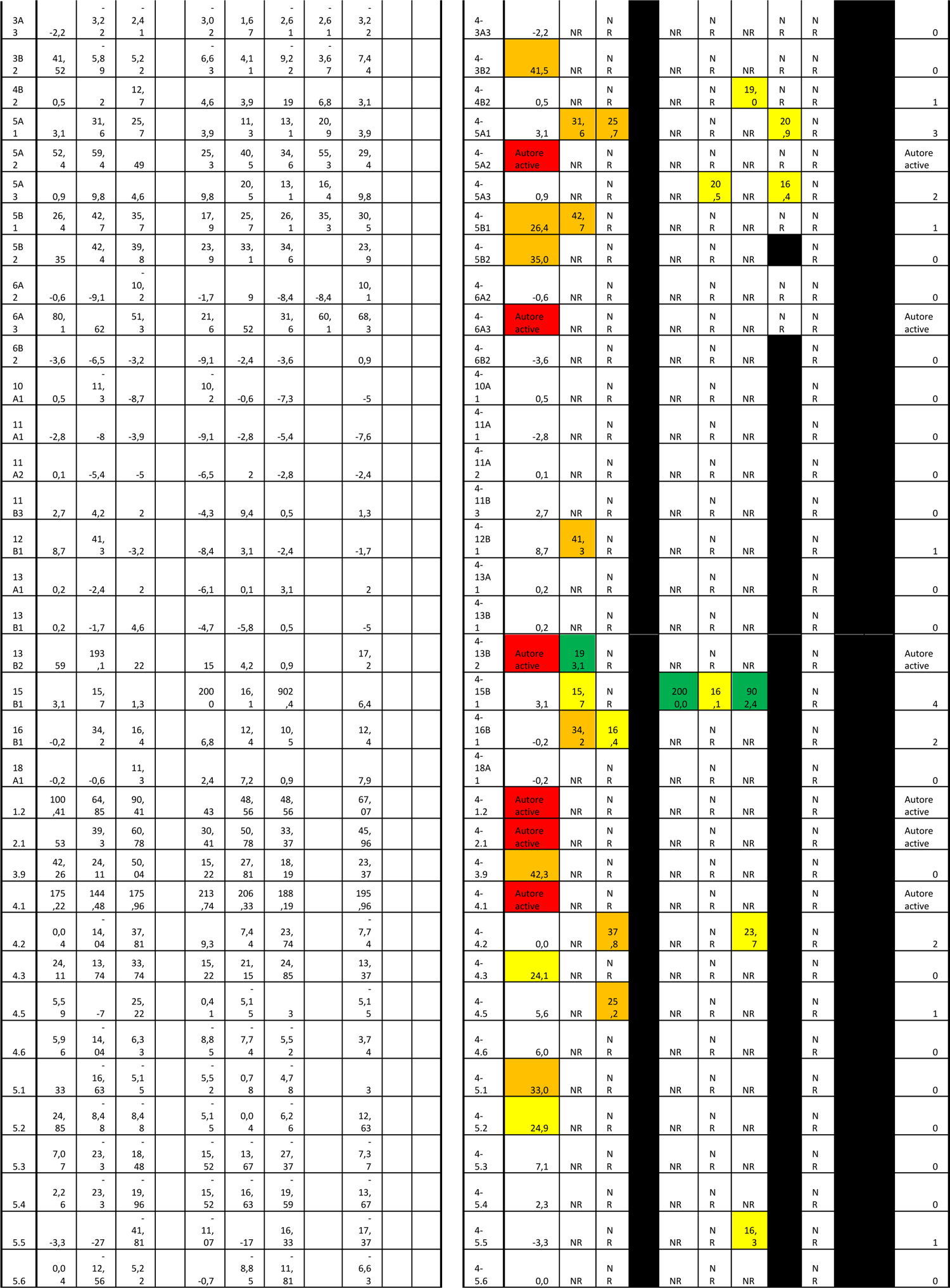
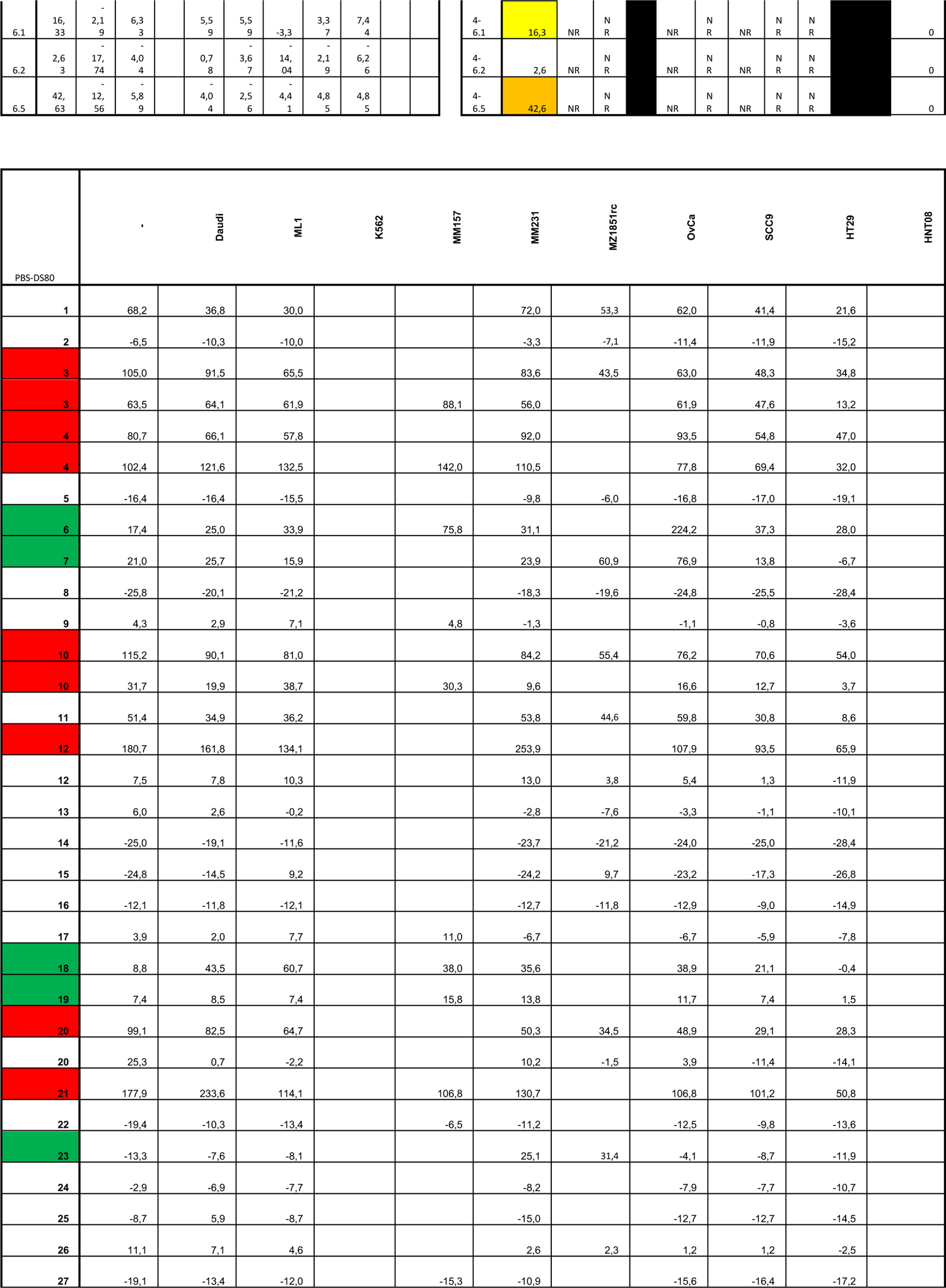
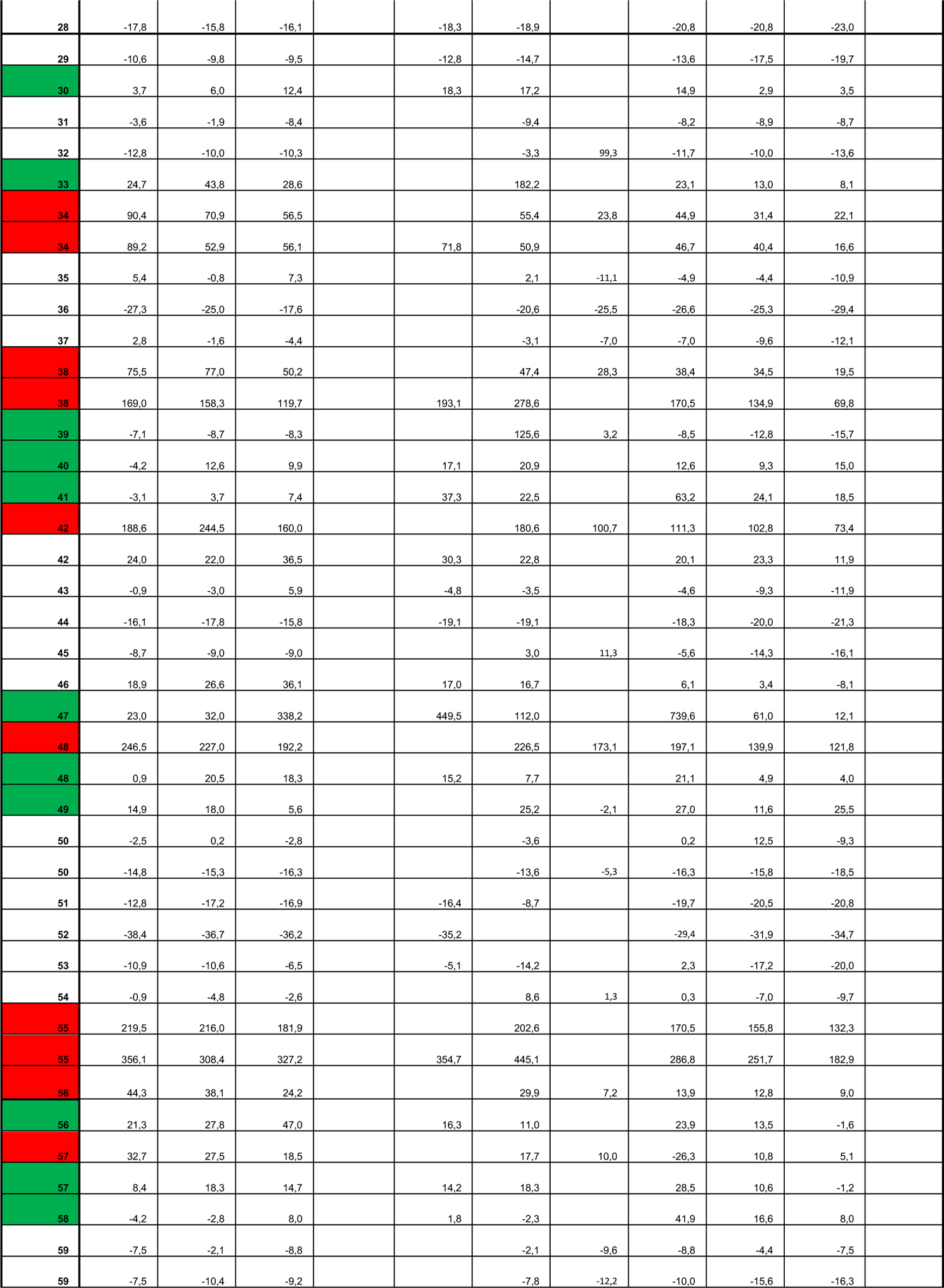
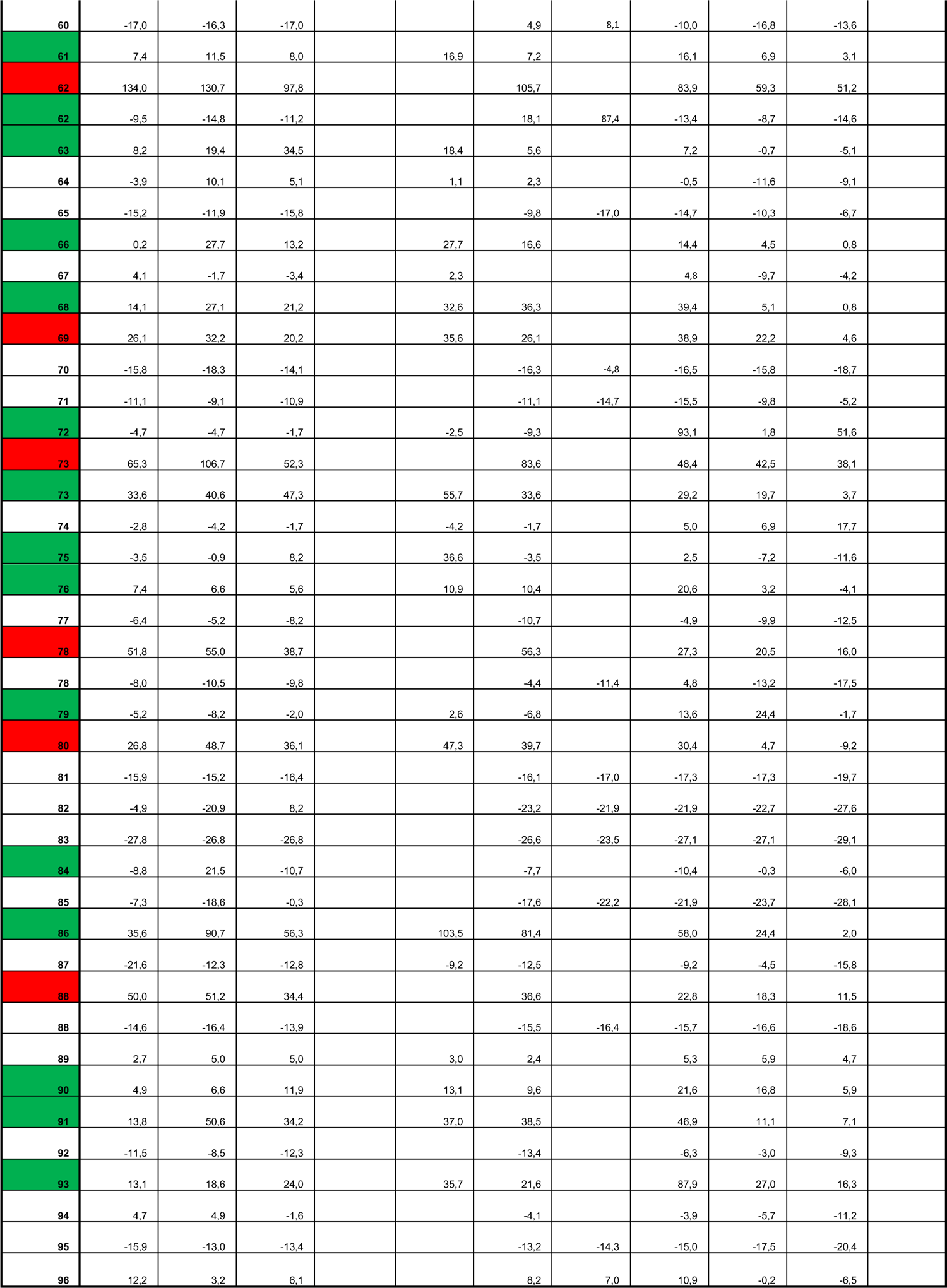
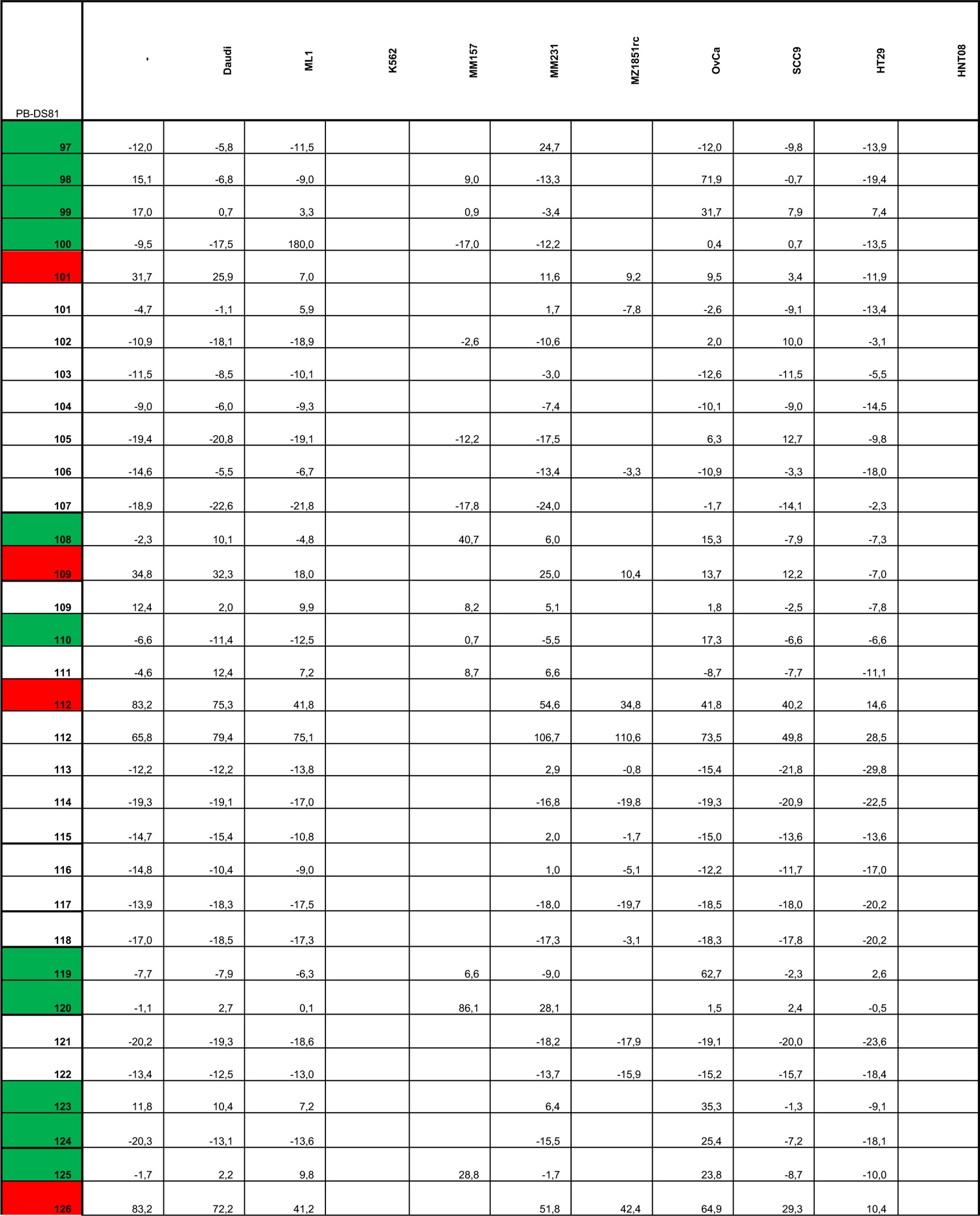
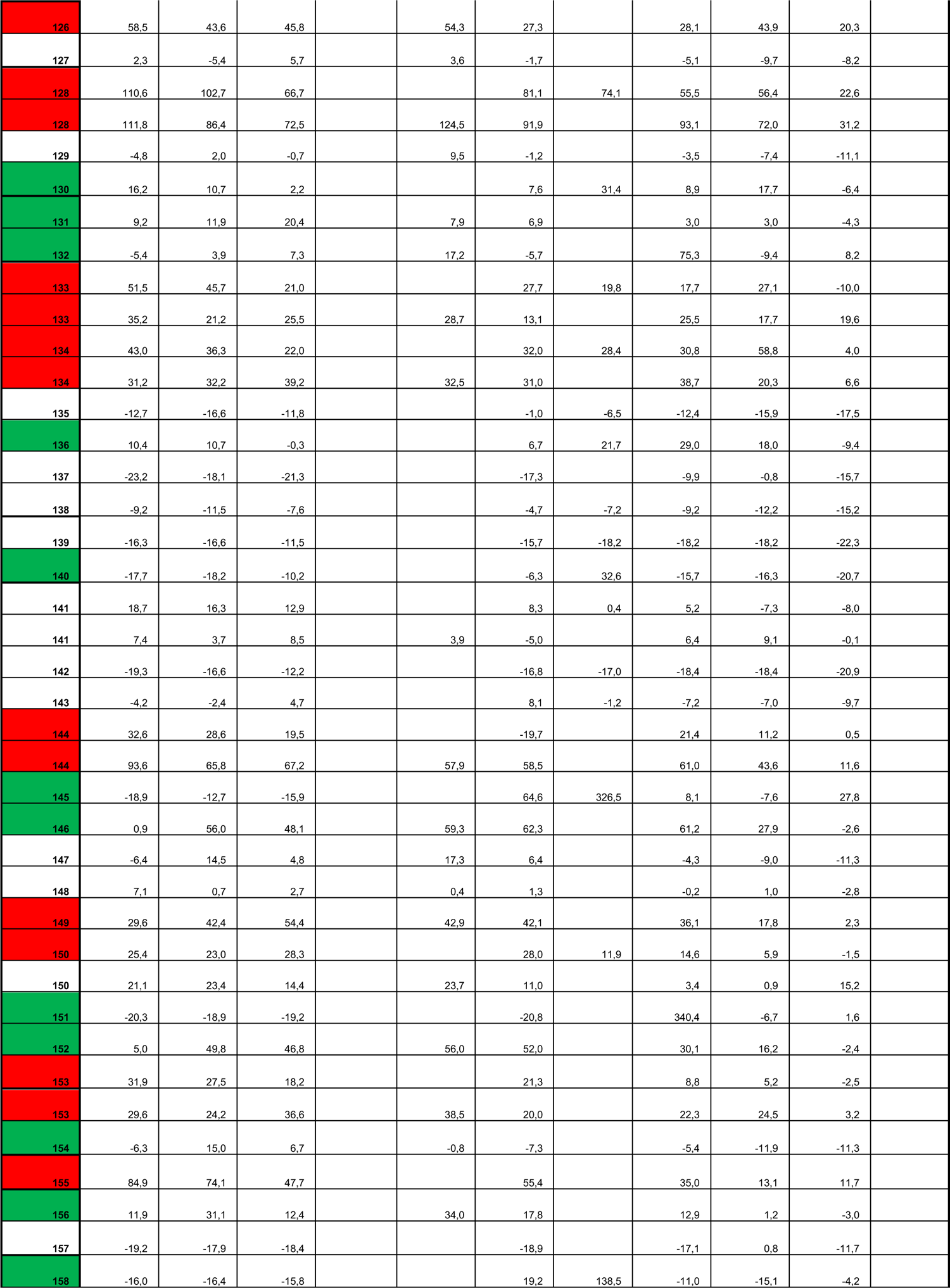
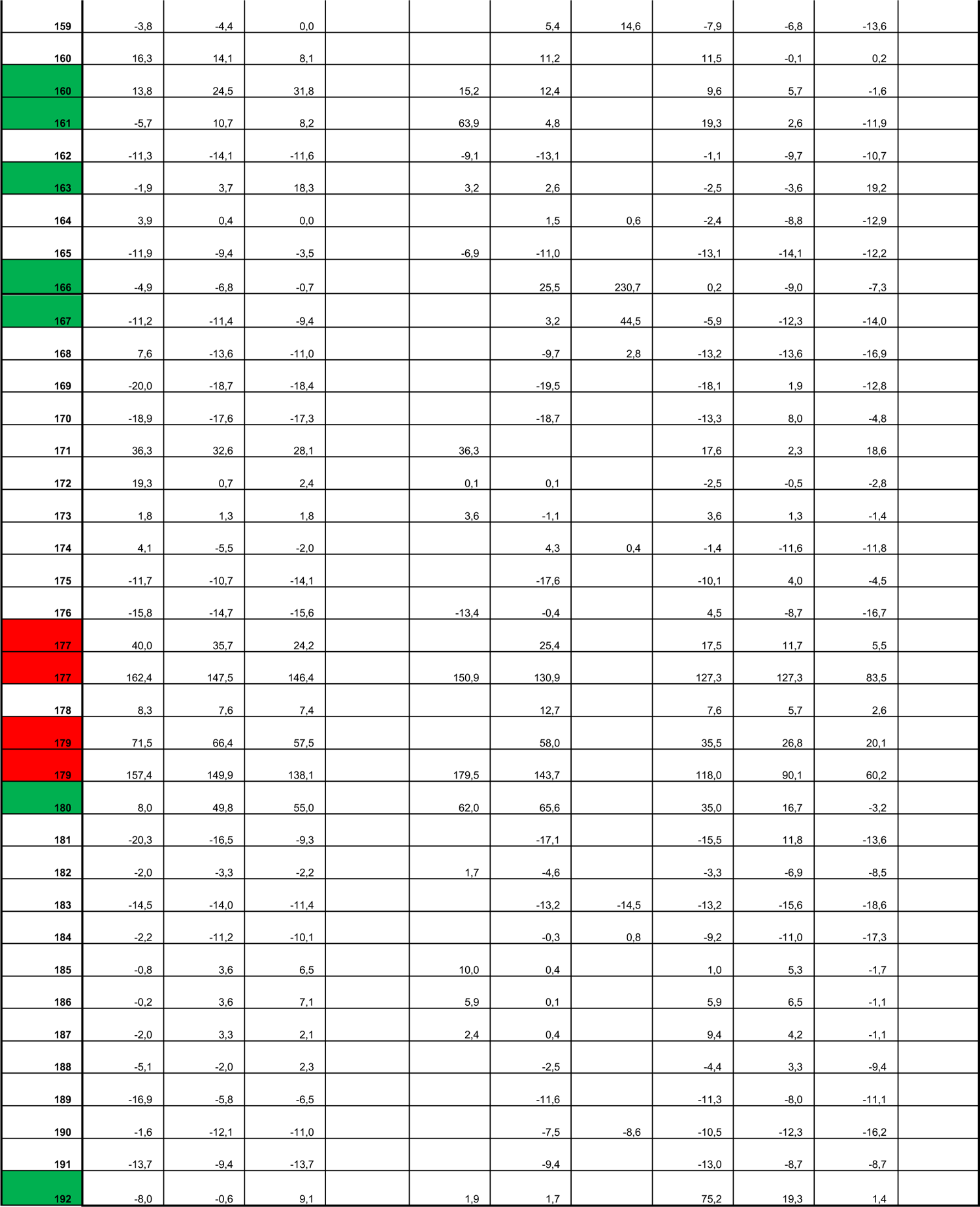
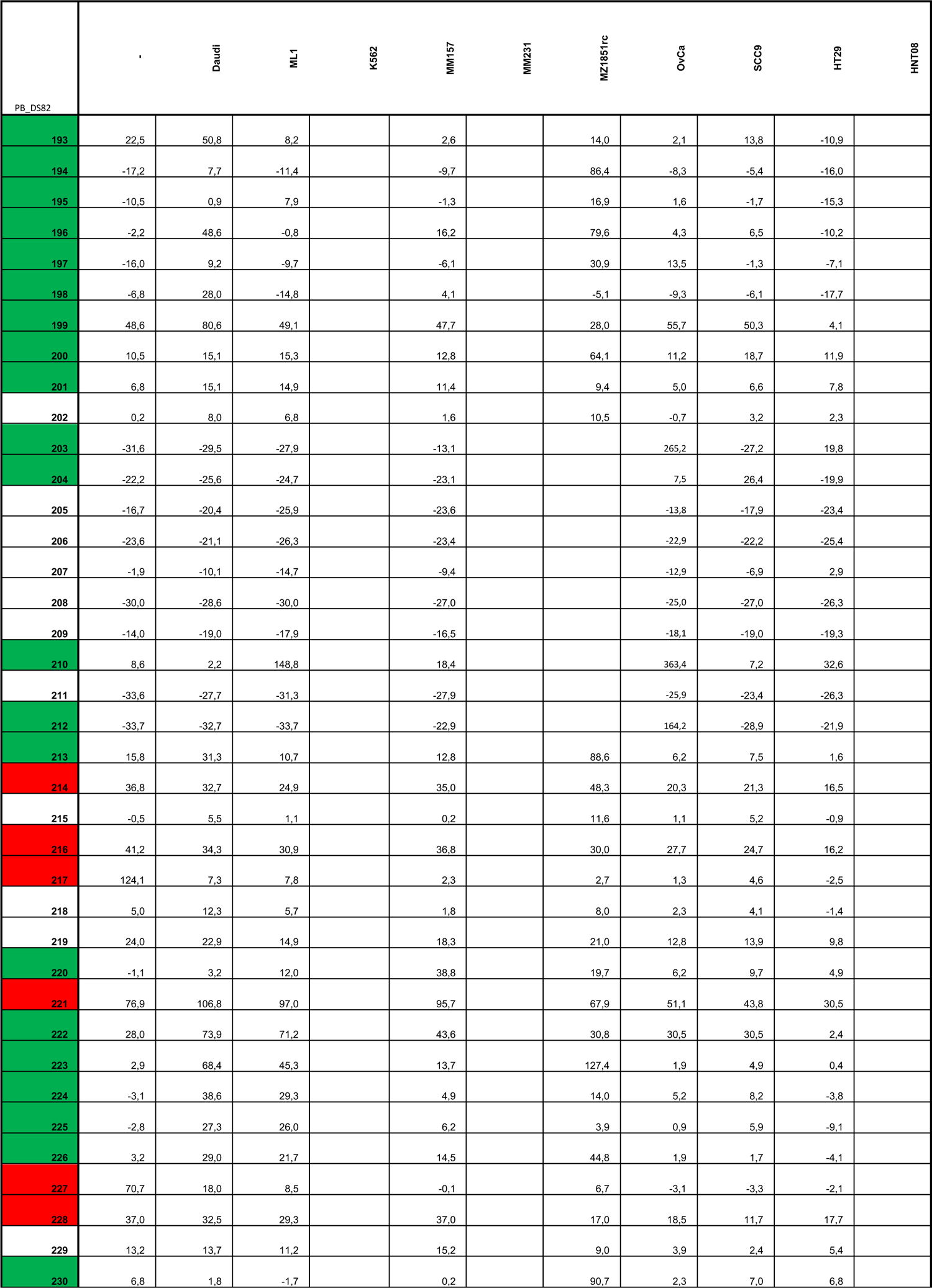
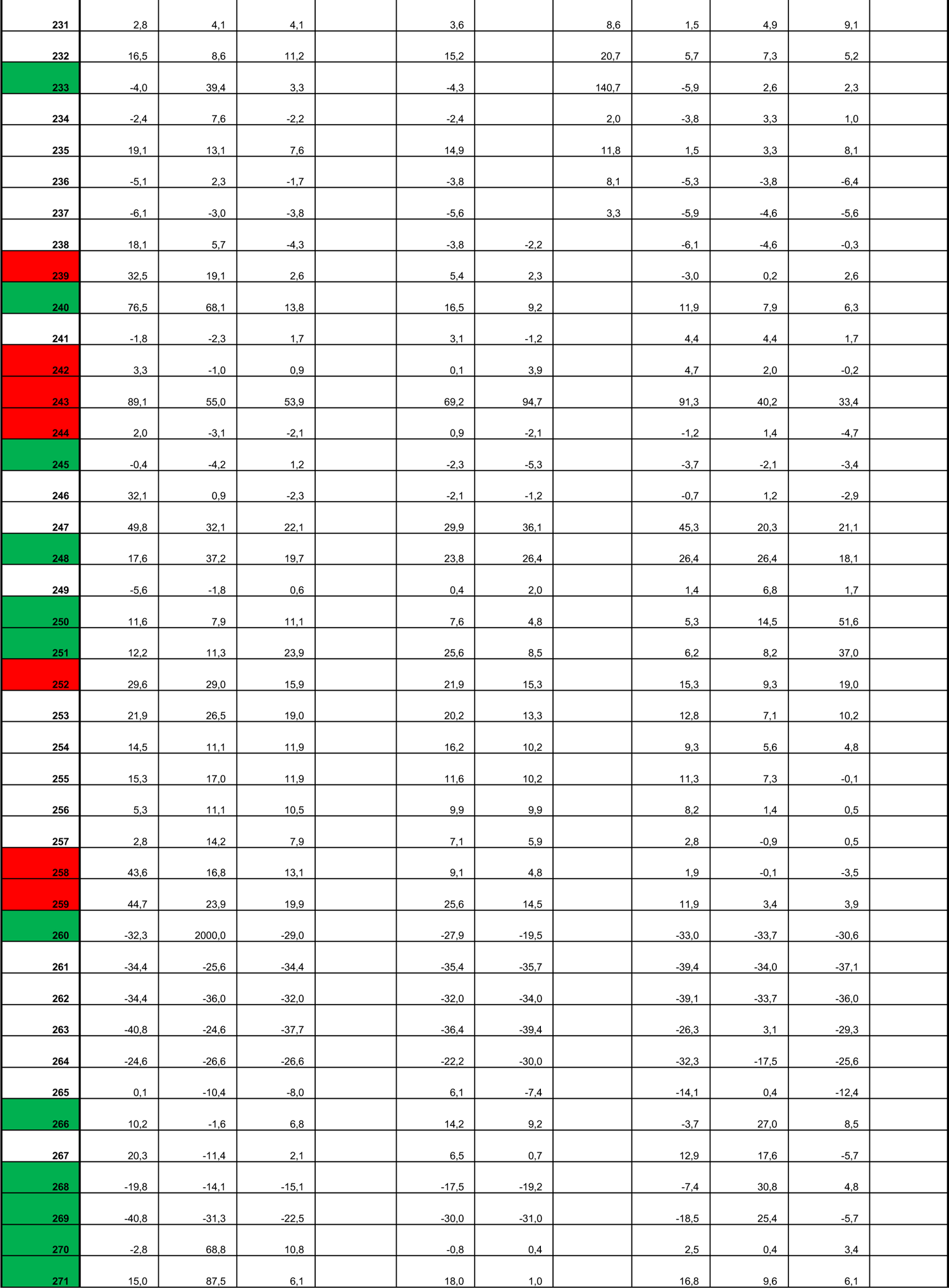
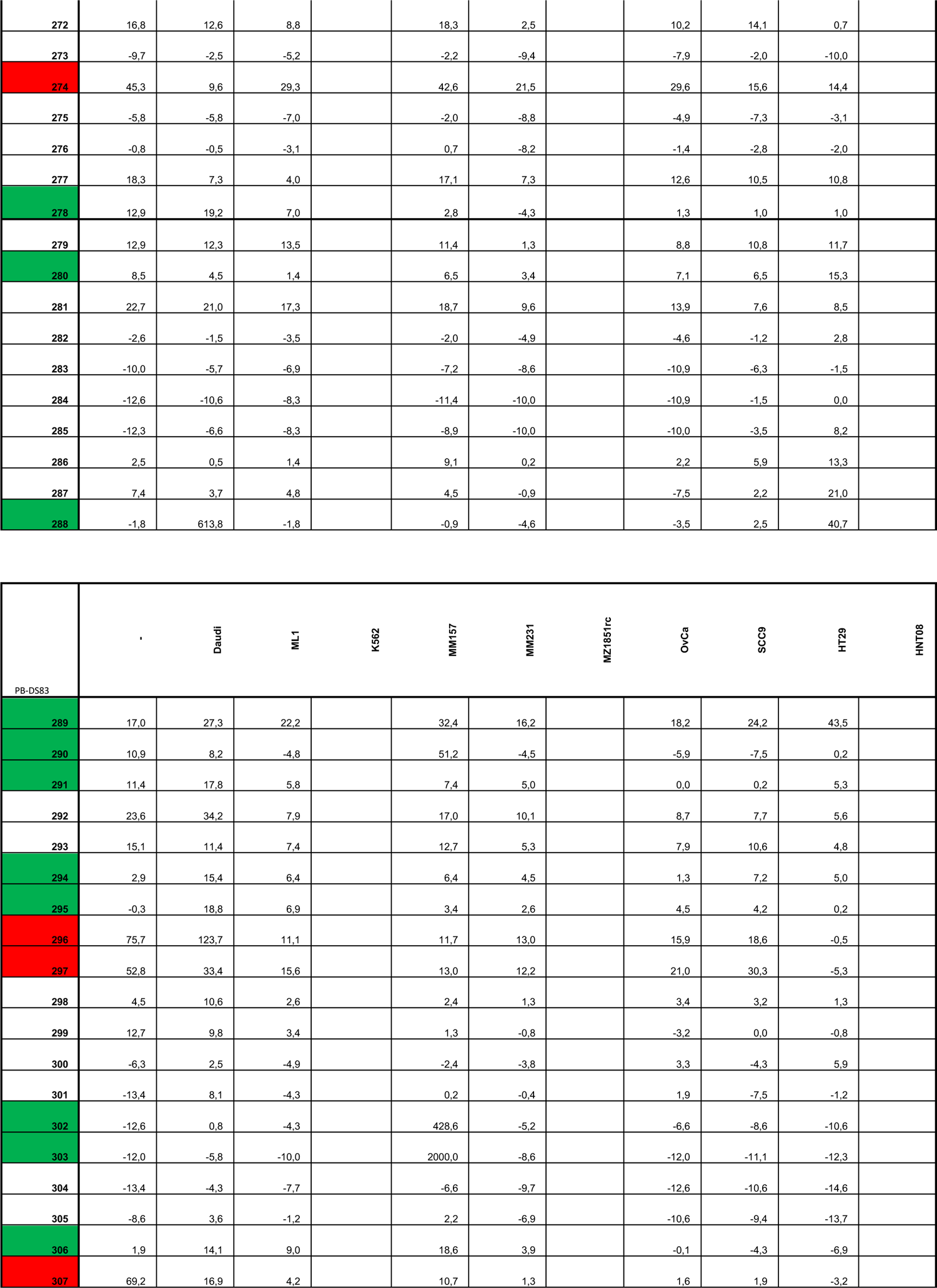
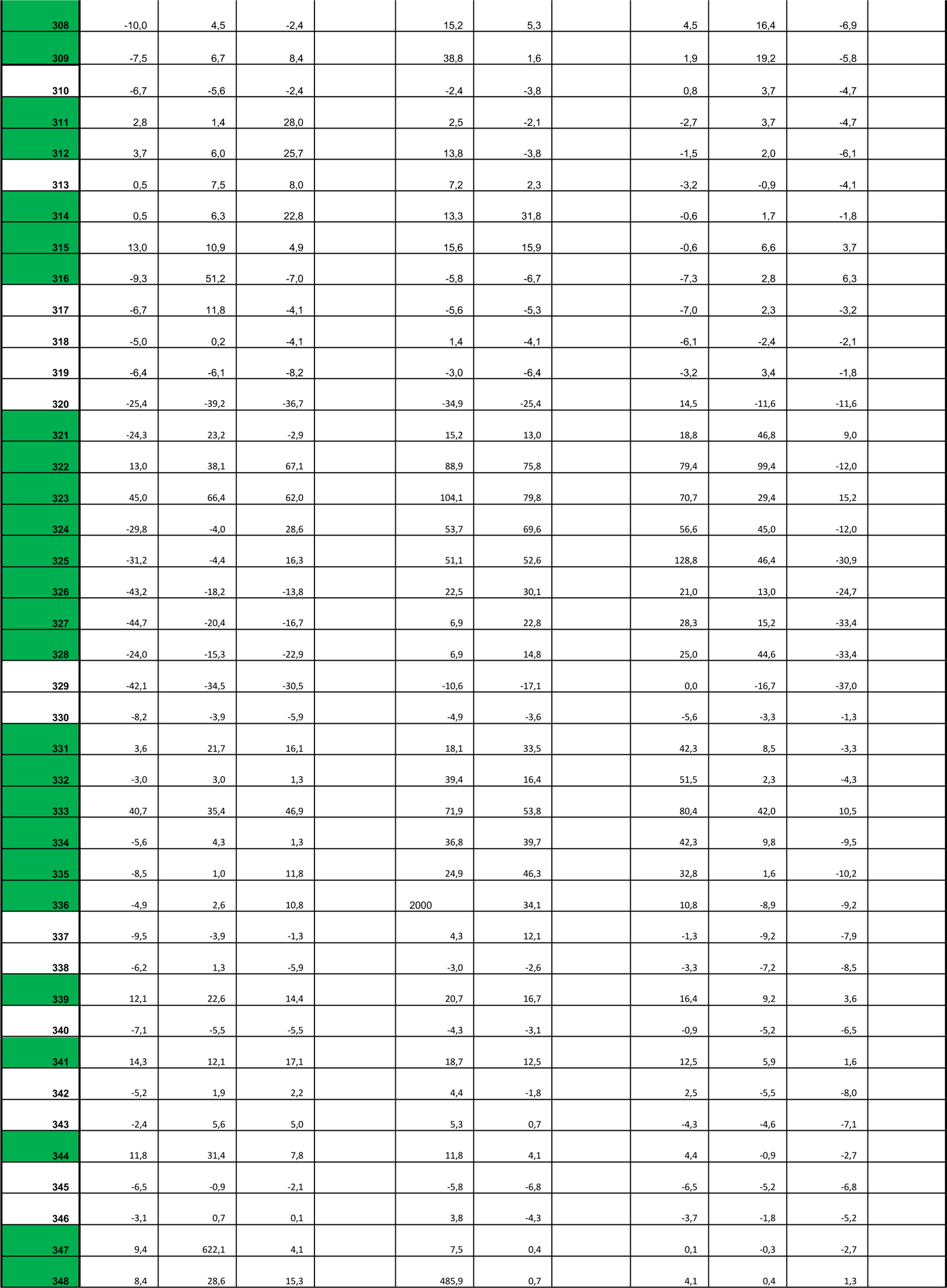
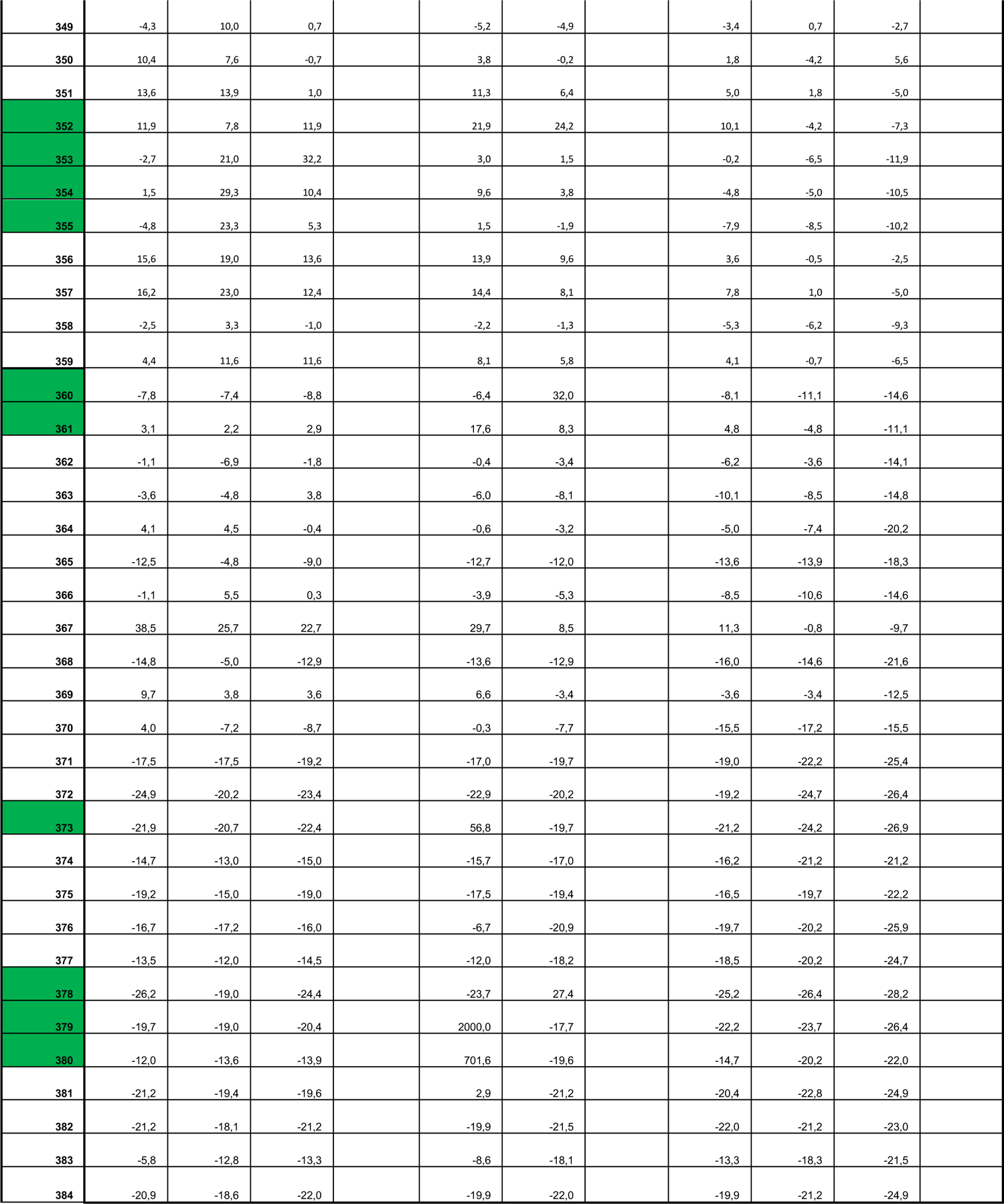
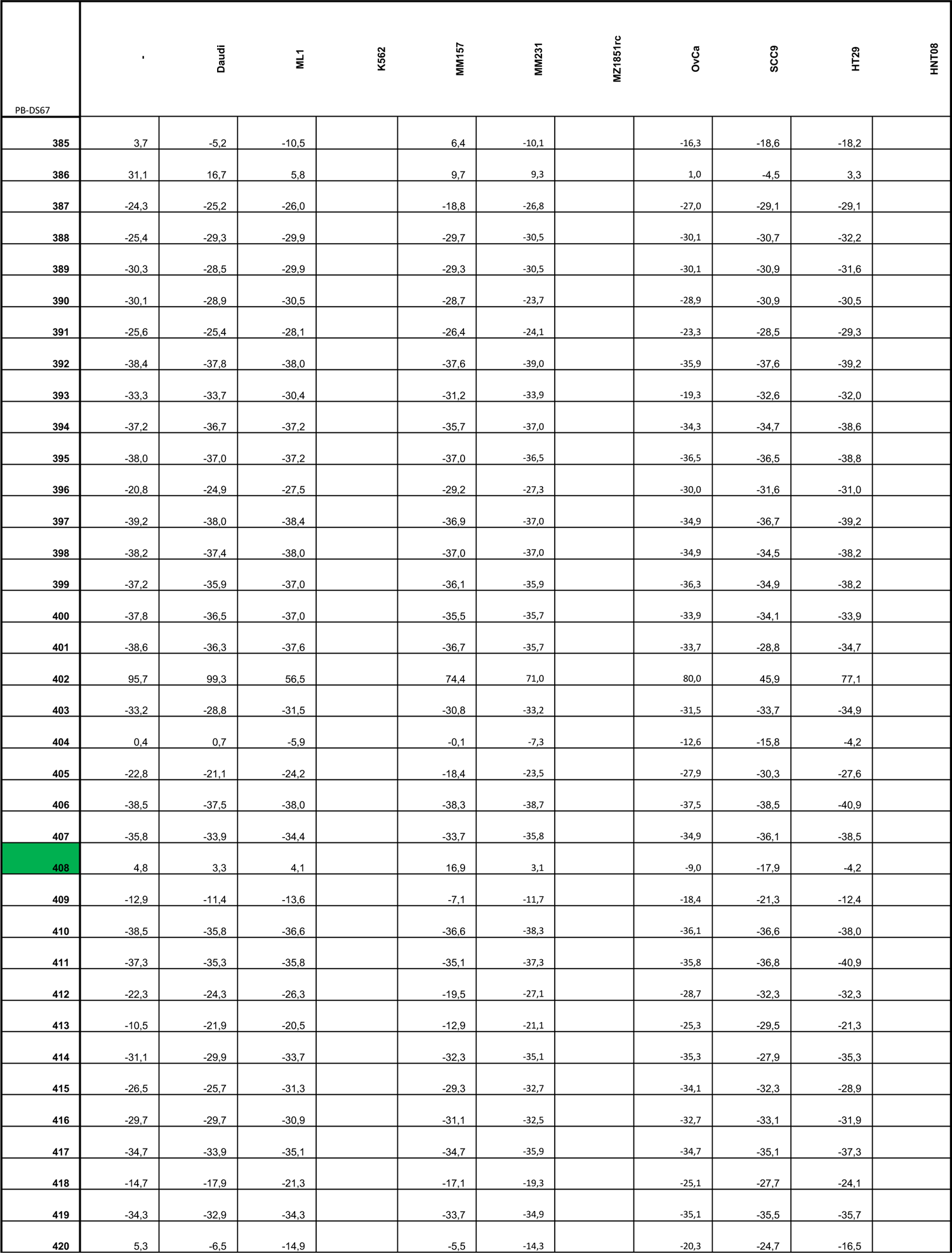
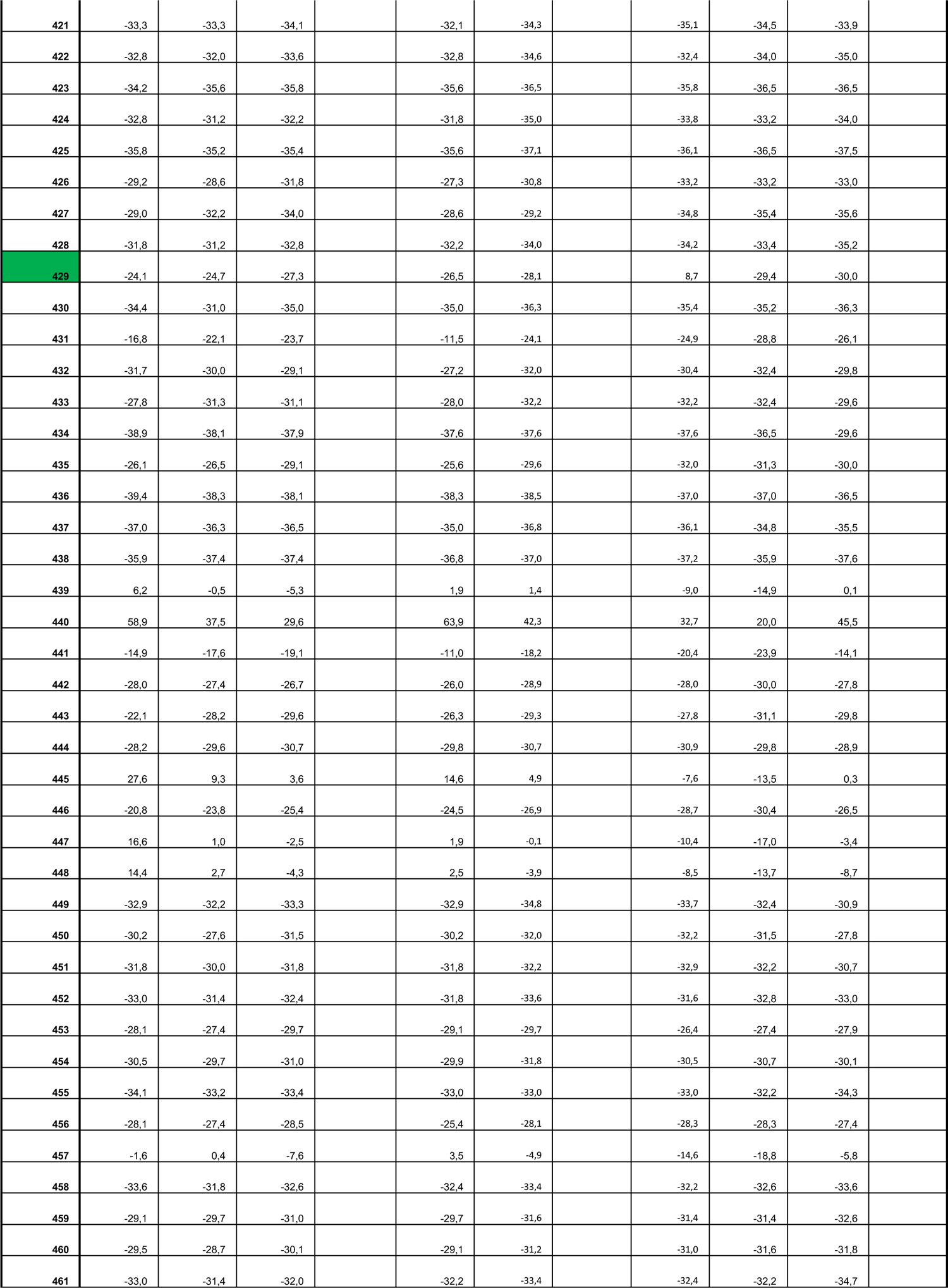
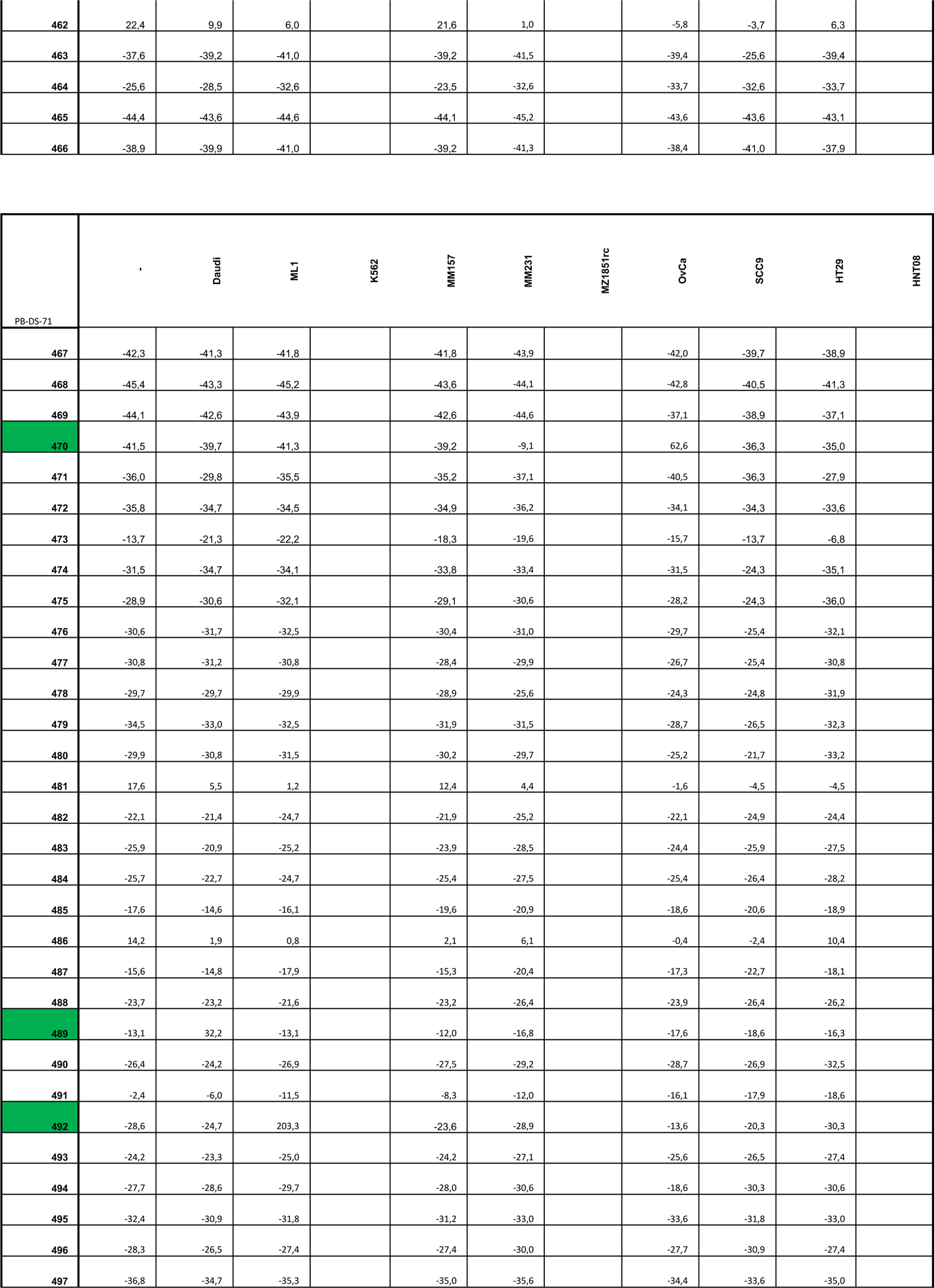
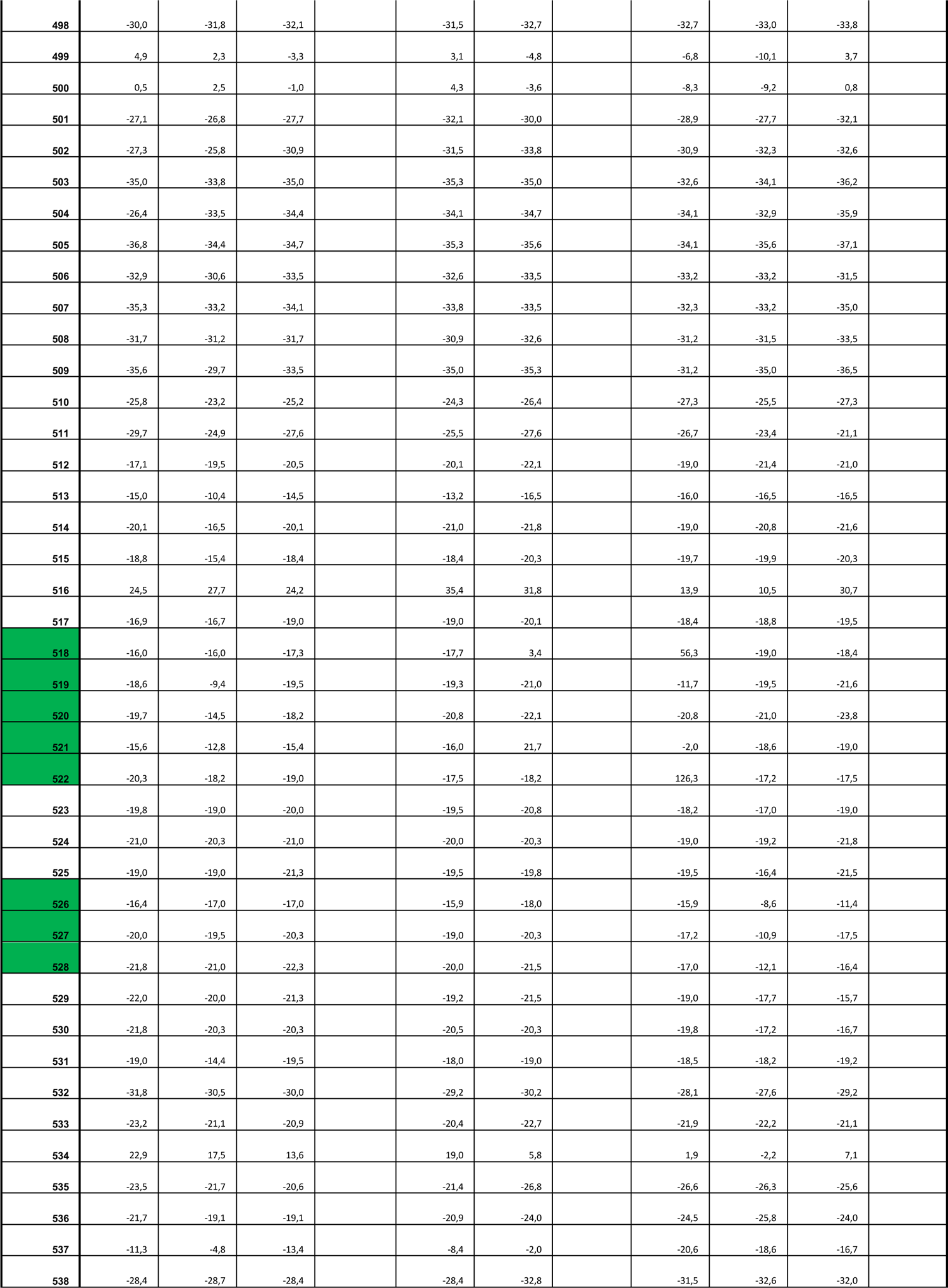
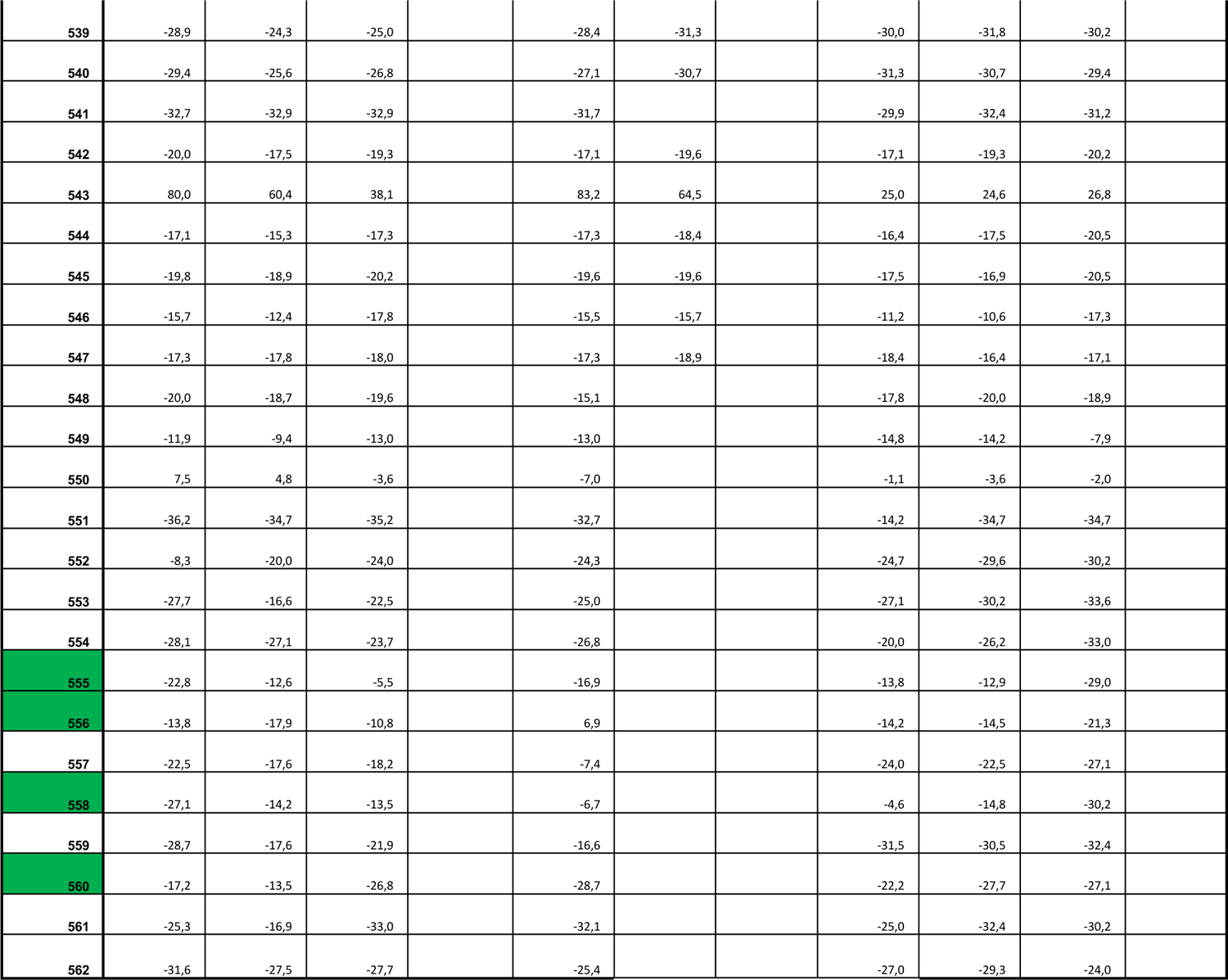

## Notes

### Competing Interest Statement

JK is shareholder of Gadeta. JK, AJ and DXB are inventors on patents gdTCR related topics.

